# Process noise distinguishes between indistinguishable population dynamics

**DOI:** 10.1101/533182

**Authors:** Matthew J. Simpson, Jacob M. Ryan, James M. McGree, Ruth E. Baker

## Abstract

*Model selection* is becoming increasingly important in mathematical biology. Model selection often involves comparing a set of observations with predictions from a suite of continuum mathematical models and selecting the model that provides the best explanation of the data. In this work we consider the more challenging problem of model selection in a stochastic setting. We consider five different stochastic models describing population growth. Through simulation we show that all five stochastic models gives rise to classical logistic growth in the limit where we consider a large number of identically prepared realisations. Therefore, comparing mean data from each of the models gives indistinguishable predictions and model selection based on population-level information is impossible. To overcome this challenge we extract *process noise* from individual realisations of each model and identify properties in the process noise that differ between the various stochastic models. Using a Bayesian framework, we show how process noise can be used successfully to make a probabilistic distinction between the various stochastic models. The relative success of this approach depends upon the identification of appropriate summary statistics and we illustrate how increasingly sophisticated summary statistics can lead to improved model selection, but this improvement comes at the cost of requiring more detailed summary statistics.

## Introduction

The logistic growth equation,

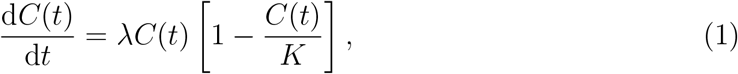

with solution,

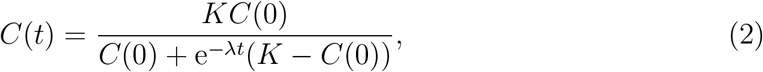

is probably the most widely used model of population dynamics in the fields of mathematical biology and mathematical ecology (Edelstein-Keshet 2005; Murray 2002). In this continuum framework, *C*(*t*) ≥ 0 is the number, or number density, of individuals, *λ* > 0 is the low-density growth rate and *K* > 0 is the carrying capacity of the environment. The key feature of the logistic growth model is that when the initial population density is small compared to the carrying capacity, *C*(0)*/K* ≪ 1, the solution takes the form of a sigmoid curve that monotonically increases from *C*(0) and asymptotes to 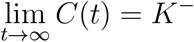.

For small initial densities, *C*(0)*/K* ≪ 1, the logistic growth model can be approximated by a simpler exponential growth model since *C*(*t*) ∼ *C*(0)e^*λt*^ for sufficiently small *t* (Warne et al. 2017). In the exponential growth model, the per capita growth rate is a positive constant, (d*C*(*t*)/d*t*)(1/*C*(*t*)) = *λ*, that is independent of the population density. This feature leads to unbounded growth as *t* increases. In contrast, the per capita growth rate for the logistic model, (d*C*(*t*)/d*t*)(1/*C*(*t*)) = *λ*[1 − *C*(*t*)/*K*], is a linearly decreasing function of *C*(*t*), such that the per capita growth rate is zero at *C*(*t*) = *K*, which leads to bounded populations at large time. This kind of transition between low-density exponential growth and high-density saturation are the two key elements of the logistic growth model that make it so appealing since these features are widely observed in many applications in biology and ecology (Edelstein-Keshet 2005; Murray 2002).

In a biological context, images in Figure 1(a)-(b) show the growth of a population of prostate cancer cells in two-dimensional cell culture at *t* = 0 and *t* = 48 h, respectively (Jin et al. 2016). This kind of growth process becomes limited by available space in the monolayer as the density increases, as illustrated in Figure 1(c). This reduced growth rate at larger densities is often modelled using Equations (1)-(2) (Browning et al. 2018; Cai et al. 2007; Sengers et al. 2007; Tremel et al. 2009). In an ecological context, the image in Figure 1(d) shows a population of polyps growing on the surface of an oyster shell. Similar to the population of cells in Figure 1(a)-(b), the net growth rate of the polyps reduces as the density increases, as illustrated in Figure 1(e). This kind of saturation can be accurately predicted using the logistic growth model (Melica et al. 2014) as illustrated in Figure 1(e). Many other kinds of ecological growth processes, such as populations of plants, are also modelled using Equations (1)-(2) (Bolker and Pacala 1997; Law et al. 2003). Far more complicated organism-level growth phenomena also exhibit saturation behaviour as illustrated by images in Figure 1(f)-(g) showing the growth of a Shiba Inu puppy at age 12 and 30 weeks, respectively. The time series showing the weight of the puppy, given in Figure 1(h), takes the form of a sigmoid curve that can be modelled using Equations (1)-(2). We note, however, that while whole organism-level growth can be approximately modelled using the classical logistic equation, as illustrated in Figure 1(h), other generalised logistic growth laws are often used to study organism-level growth (Frascoli et al. 2014; Goriely 2017; Tsoularis and Wallace 2002).

**Figure 1:**
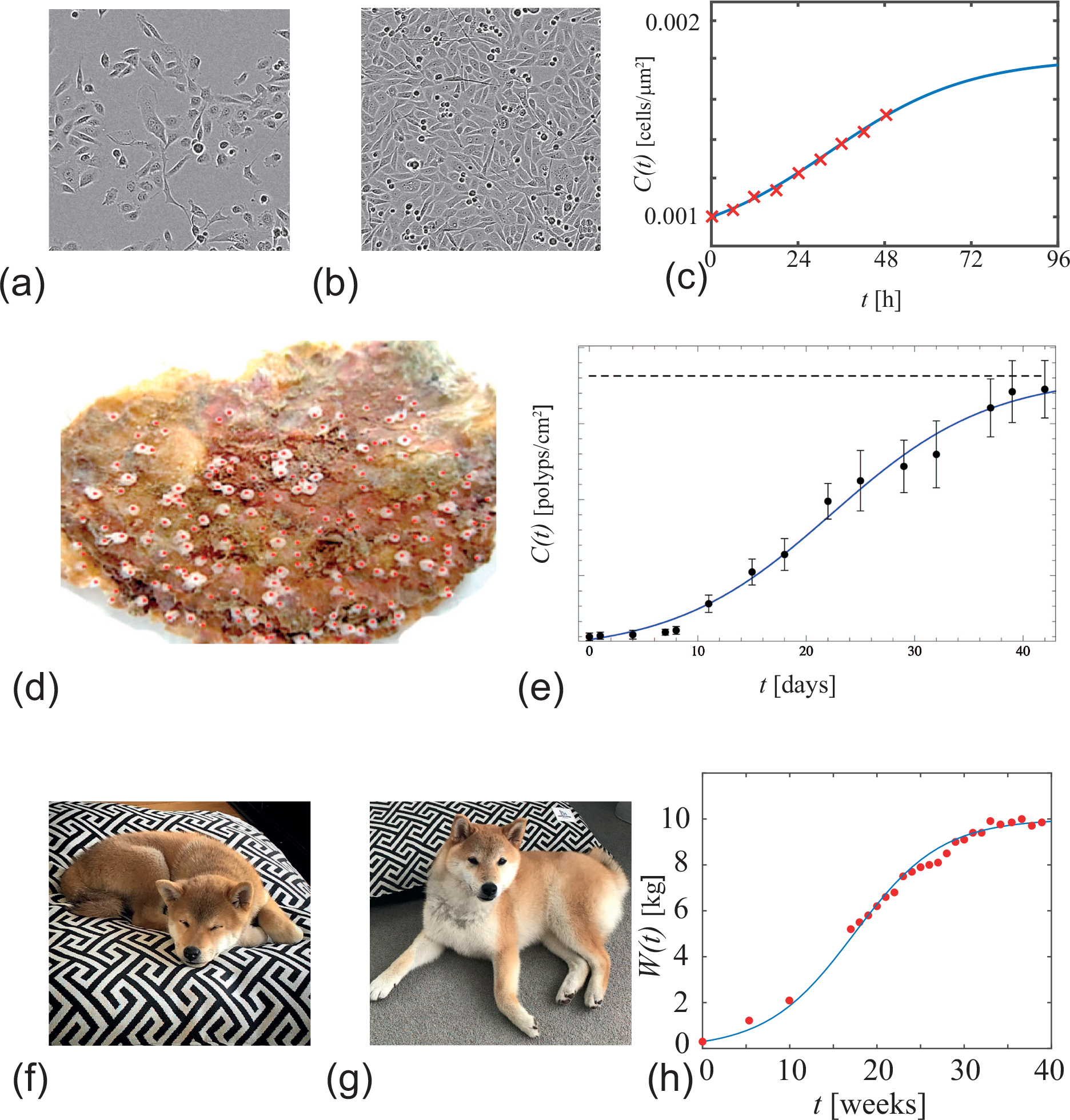
Examples of logistic growth-type phenomena. (a)-(b) shows images of a cell proliferation assay performed with prostate cancer cells [23]. The images in (a) and (b) correspond to *t* = 0 and *t* = 48 h, respectively, and each image shows a region that is approximately 450 *μ*m^2^ in area. The plot in (c) shows experimental estimates of cell density, *C*(*t*), (red crosses) superimposed on a plot of Equation (2) with *C*(0) = 0.001 cells/*μ*m^2^, *K* = 0.002 cells/*μ*m^2^ and *λ* = 0.04 /h. (d) shows an oyster shell, upon which a population of polys (red) are growing. The plot in (e) shows experimental estimates of the polyp density, *C*(*t*), superimposed on a plot of Equation (2) with *C*(0) = 0.081 polyps/cm^2^, *K* = 1.81 polyps/cm^2^ and *λ* = 0.006 /h. Images in (f) and (g) show photographs of a Shiba Inu puppy *Frankie*, aged 12 and 35 weeks, respectively. Data in (h) show time series measurements of Frankie’s weight, *w*(*t*) (red dots) superimposed on a plot of Equation (2) with *C*(0) = 0.35 kg, *K* = 10.7 kg and *λ* = 0.00113 /h. Images and data in (a)-(c) are reproduced with permission from Jin et al. (2016). Images and data in (d)-(e) are reproduced with permission from Melica et al. (2014). Images in (f) and (g), and data in (h) were obtained by the first author.

Instead of working with a continuum-level description of population dynamics, such as Equations (1)-(2), it is often the case in mathematical biology and mathematical ecology that stochastic discrete models are used (Bolker and Pacala, 1997; Codling et al. 2008; Ermentrout and Edelstein-Keshet 1993; Lambert et al. 2018; Law et al. 2003; Maclaren et al. 2015). Individual-based stochastic models are sometimes preferred over continuum models since continuum models can not provide information about individual-level behaviour and continuum models do not capture or predict stochastically (Frascoli et al. 2013; Treloar et al. 2014; O’Dea and King 2012).

In this section we describe five different discrete models of population dynamics and explain their relationship to Equations (1)-(2). In our description of these models we pay careful attention to illustrate the different mechanisms inherent in each model, and we describe how each model can be simulated to produce stochastic realisations. Using results of these stochastic realisations, we demonstrate that each of the different discrete models gives rise to indistinguishable logistic growth in the limit of a very large number of identically prepared realisations. Therefore, when we are dealing with noisy individual realisations of the discrete models we are considering random variables. In contrast, when we consider the averaged outcome of a large number of identically prepared realisations of the discrete models we are dealing with deterministic variables. To make this distinction clear we denote the number of individuals in any realisation from any one of the five discrete models as 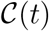, and the average number of individuals constructed using a largenumber of identically prepared realisations as

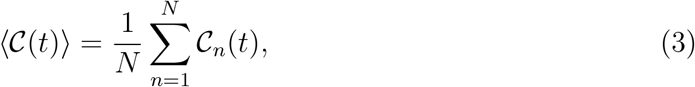

where 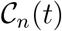 is the number individuals in the *n*^th^ identically prepared realisation of a particular stochastic model at time *t*, and *N* is the total number of realisations used to construct the averaged density.

## Discrete models

We will now describe and implement the five completely distinct stochastic models that we consider in this work. The first three models are *spatially implicit* models meaning that we consider the population of individuals as being spatially well-mixed but we do not explicitly consider the spatial location of any particular individual (Geritz and Kisdi, 2012). In contrast, the last two models are *spatially explicit* and we explicitly track the position of each individual within the population during the simulations (Baker and Simpson, 2010; Simpson et al. 2010). In the spatially explicit models we initially enforce the individuals in the population to be well mixed. At the beginning of each simulation, individuals are placed uniformly, at random, taking care to ensure that no two individuals occupy the same location. Each of the five discrete models will be simulated using the Gillespie algorithm (Gillespie, 1977), and we will now describe these models and the key variables associated with each model.

### 2.1 Model 1: Spatially implicit birth-only

The first model considers a population of individuals, 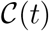, that undergoes a birth process where the net birth rate is a linearly-decreasing function of the total density. This gives rise to a per capita growth rate of

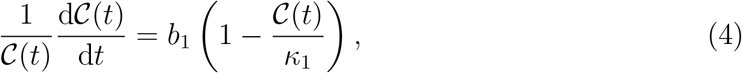

where *b*_1_ > 0 is the low density birth rate and *κ*_1_ > 0 is the carrying capacity of the environment (Geritz and Kisdi, 2012).

### 2.2 Model 2: Spatially implicit birth-death

The second model considers a population of individuals, 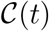, that undergoes a combined birth-death process. Here, the net birth rate is a linearly-decreasing function of the total density and the death rate is constant. This gives rise to a per capita growth rate of

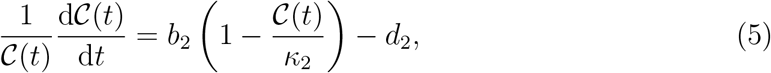

where *b*_2_ > 0 is the low density birth rate, *d*_2_ > 0 is the death rate, and *κ*_2_ > 0 is the carrying capacity of the environment (Geritz and Kisdi, 2012). Since we are interested in populations that grow, as opposed to populations that become extinct, we will focus on parameter choices where *d*_2_/*b*_2_ ≪ 1.

### 2.3 Model 3: Spatially implicit birth-death-annihilation

The third model considers a population of individuals, 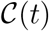, that undergoes a combination of birth, death and annihilation processes. Here, the birth and death rates are constants, and the annihilation mechanism can be thought of representing the case where two individuals compete with each other for some kind of resource, leading to the destruction of one of the individuals. Together, these three mechanisms give rise to a per capita growth rate of

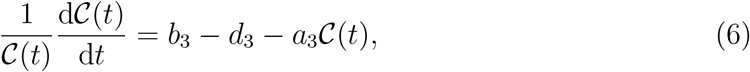

where *b*_3_ > 0 is birth rate, *d*_3_ > 0 is the death rate and *a*_3_ > 0 is the rate of loss due to annihilation (Geritz and Kisdi, 2012). Again, since we are interested in the case of growing populations we will focus on parameter choices where *d*_3_/*b*_3_ ≪ 1 and *a*_3_/*b*_3_ ≪ 1.

### 2.4 Model 4: Spatially explicit birth-only

Unlike the first three models where we do not consider the spatial location of individual agents in the population, we now consider a spatially explicit model where individuals within the population reside on a three-dimensional square lattice, with lattice spacing Δ, and the domain is a cube with side length *L* so that the total number of lattice sites is (*L/*Δ)^3^. All simulations involve periodic boundary conditions and the model is an *exclusion process* meaning that each lattice site can be occupied by, at most, a single individual (Liggett, 1999; Simpson et al. 2010). All simulations are initialised by choosing a particular number of individuals, 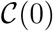, and distributing them uniformly, at random, across the lattice. Each time we initialise the model we take care to ensure that each site is occupied by no more than a single agent. During the simulations agents undergo one of two types of events. First, agents undergo movement events, at rate *m*_4_ > 0 (Baker and Simpson, 2010). During a potential movement event, the agent will attempt to move to one of the six nearest neighbour lattice sites with equal probability. If the chosen target site is occupied then that potential event will be aborted. Second, agents undergo birth events, at rate *b*_4_ > 0. During a potential birth event the mother agent will attempt to place a daughter agent on one of the six nearest neighbour lattice sites with equal probability. If those chosen target site is occupied then that potential proliferation event is aborted (Baker and Simpson, 2010).

Previously, Baker and Simpson (2010) explain how to construct the continuum limit of this model and show that, for this well-mixed (translationally invariant) initial condition the continuum limit, written in terms of the per capita growth rate, can be written as

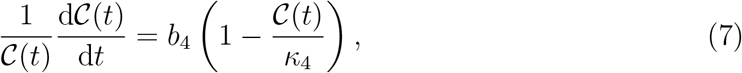

where *b*_4_ > 0 is birth rate and *κ*_4_ = (*L/*Δ)^3^ is the total number of lattice sites. We note that while the movement rate, *m*_4_, does not appear explicitly in the continuum limit description, we have the additional requirement that *b*_4_/*m*_4_ ≪ 1 for the continuum limit description to hold, and more details about this condition are given by Baker and Simpson (2010).

### 2.5 Model 5: Spatially explicit birth-death

The fifth and final model we consider is an extension of the spatially explicit birth only model, except that now we consider both birth and death events. Again, we consider a spatially explicit model where individuals within the population reside on a three-dimensional square lattice, with lattice spacing Δ, and the domain is a cube with side length *L* so that the total number of lattice sites is (*L/*Δ)^3^. All simulations involve periodic boundary conditions and the model is an exclusion process meaning that each lattice site can be occupied by, at most, a single individual. All simulations are initialised by choosing a particular number of individuals, 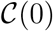, and randomly distributing them across the lattice, taking care to ensure that each site is occupied by no more than a single agent. During the simulations agents undergo three types of events. Firstly, agents undergo movement events, at rate *m*_5_ > 0, in exactly the same way as for model 4. Second, agents undergo birth events, at rate *b*_5_ > 0, in exactly the same way as for model 4. Finally, agents undergo death events, at rate *d*_5_ > 0, where agents are simply removed from the lattice.

Again, Baker and Simpson (2010) explain how to construct the continuum limit of this model and show that for this translationally invariant initial condition the continuum limit, written in terms of the per capita growth rate, is given by

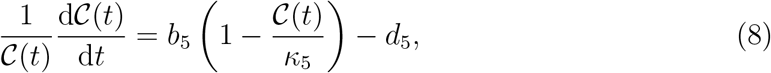

where *b*_5_ > 0 is birth rate, *d*_5_ > 0 is the death rate, and *κ*_5_ = (*L/*Δ)^3^ is the total number of lattice sites. Again, while the movement rate, *m*_5_, does not appear explicitly in the continuum limit description, we have the additional requirement that *b*_5_/*m*_5_ ≪ 1 and *d*_5_/*m*_5_ ≪ 1 for the continuum limit description to hold (Baker and Simpson, 2010). Furthermore, since we are interested in population growth, we focus on parameter choices with *d*_5_/*b*_5_ ≪ 1.

## 3 Results and Discussion

### 3.1 Re-scaling and population-level indistinguishability

Each of the models 1-5 are now simulated using the Gillespie algorithm, and in each case we always observe that the population evolves from 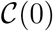 to approach some long-time positive steady state population density 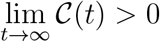. Some re-scaling of the parameters in Equations (4)-(8), according to the relationships summarised in Table 1, suggests that despite the major individual-level differences in each of the five discrete models, the average behaviour we expect to see from all five stochastic models is neatly described by Equations (1)-(2).

**Table 1:**
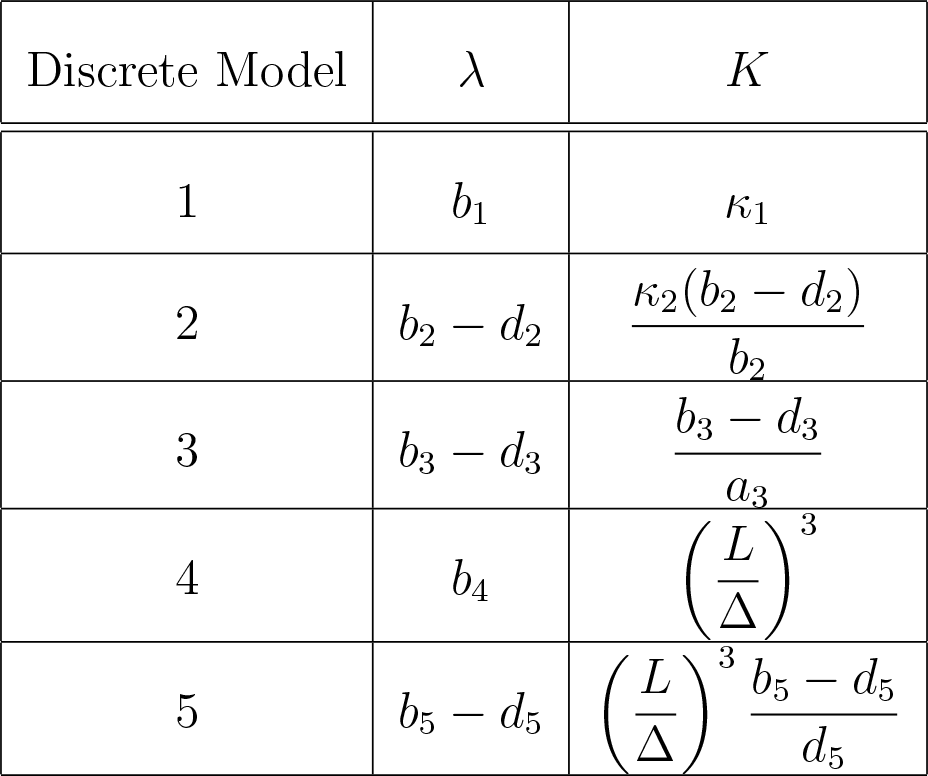
Re-scaling of the parameters in each discrete model.

With this framework we are now in a position to select biologically-relevant parameter values in Equations (1)-(2) and perform a suite of simulations from the five discrete models, using the equivalent parameters in Table 1, and make a comparison of the predictions of the stochastic and continuum models. To select the parameter values in Equations (1)-(2) we focus on population growth in the context of biological cells (Browning et al. 2018; Cai et al. 2007; Sengers et al. 2007). We note that a typical *in vivo* cell doubling time is approximately 20-30 h, giving an estimate of *λ* = 0.02 − 0.03 /h (Jin et al. 2016). In contrast, two-dimensional *in vitro* experiments, with plentiful nutrients and oxygen, are associated with a much faster doubling time of approximately 14 h, giving an estimate of *λ* = 0.05 /h (Jin et al. 2016). Therefore, in this work we will consider biologically relevant upper and lower bounds on the value of *λ*. In the main document we consider results for *λ* = 0.01 /h and we present a second set of results for *λ* = 0.05 /h in the Supplementary Material document. In all cases we set *C*(0) = 100 and *K* = 1000 so that we consider net population growth of almost an order of magnitude. Furthermore, when we present results graphically we always plot 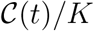 and *C*(*t*)/*K* as a function of time so that the results are presented in terms of population densities relative to the carrying capacity.

Results in Figure 2 show single realisations of each stochastic model and we compare the evolution of 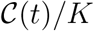 from each realisation with *C*(*t*)/*K* from Equations (1)-(2). In each case we see that stochastic models are noisy but the agreement with the continuum solution is clear. This match between the simulation data and the solution of the continuum model suggests that if we are provided with data showing 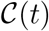 or 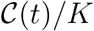, as in Figure 1(c), (e) and (h), it would be very difficult to distinguish between which of the five discrete models provides the best explanation of the experimental data.

**Figure 2:**
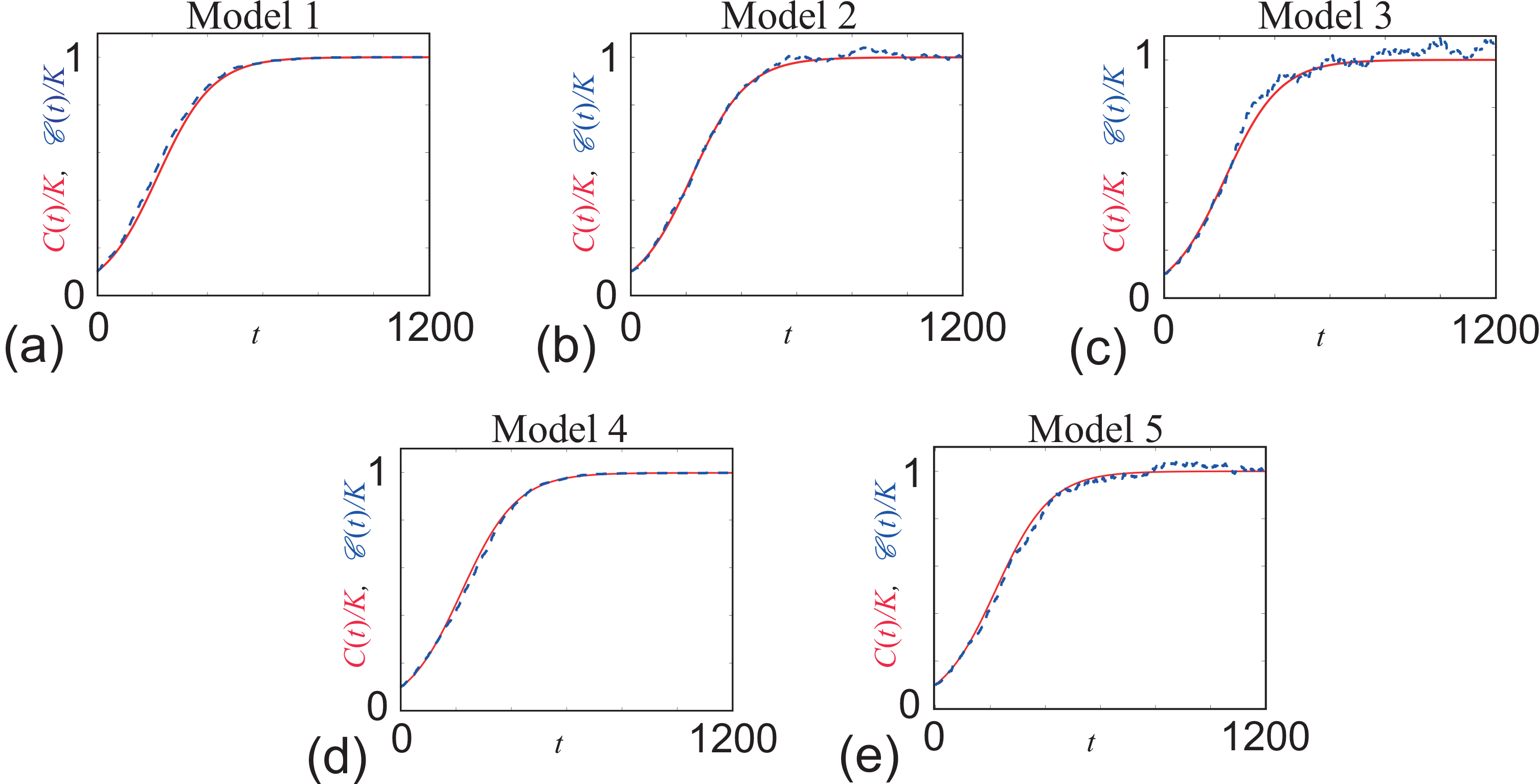
Comparison of the solution of the classical logistic equation, 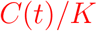, with single realisations of the five different stochastic models, 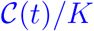. The continuum-discrete match for models 1-5 are shown in (a)-(e), respectively, with 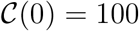, *K* = 1000 and *λ* = 0.01 /h. Parameters in the five discrete models are: *b*_1_ = 0.01 /h, *κ*_1_ = 1000; *b*_2_ = 0.0125 /h, *d*_2_ = 0.0025 /h, *κ*_2_ = 1250; *b*_3_ = 0.0125 /h, *d*_3_ = 0.0025 /h, *a*_3_ = 0.00001 agents/h; *L* = 10, Δ = 1, *b*_4_ = 0.01 /h, *m*_4_ = 1 /h; *L* = 10, Δ = 1, *b*_5_ = 0.0125 /h, *d*_5_ = 0.0025 /h, *m*_5_ = 1 /h.

The challenge of model selection becomes far more acute if we take the standard approach to interpret noisy data from a stochastic model and consider averaging the results from the discrete model over many identically prepared realisations. In any single realisation of any of the discrete models, events occur at random times and the time between events is exponentially distributed (Gillespie, 1977). To construct averaged density profiles we first interpolate the discrete time population data to give a continuous description of 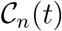. To interpolate the discrete time data we use MATLABs `previous` interpolation scheme (Mathworks 2019a) to give continuous representations of 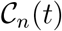 for 0 ≤ *t* ≤ 1200. We then construct averaged density profiles by averaging the values of 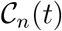 at 1201 equally spaced times, *t* = 0, 1, 2, …, 1200, according to Equation (3). Results in Figure 3 compare the solution of Equations (1)-(2) with averaged data from each discrete model, 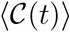, with *N* = 1000. In this case we see that *C*(*t*) and 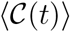 are visually indistinguishable at this scale.

**Figure 3:**
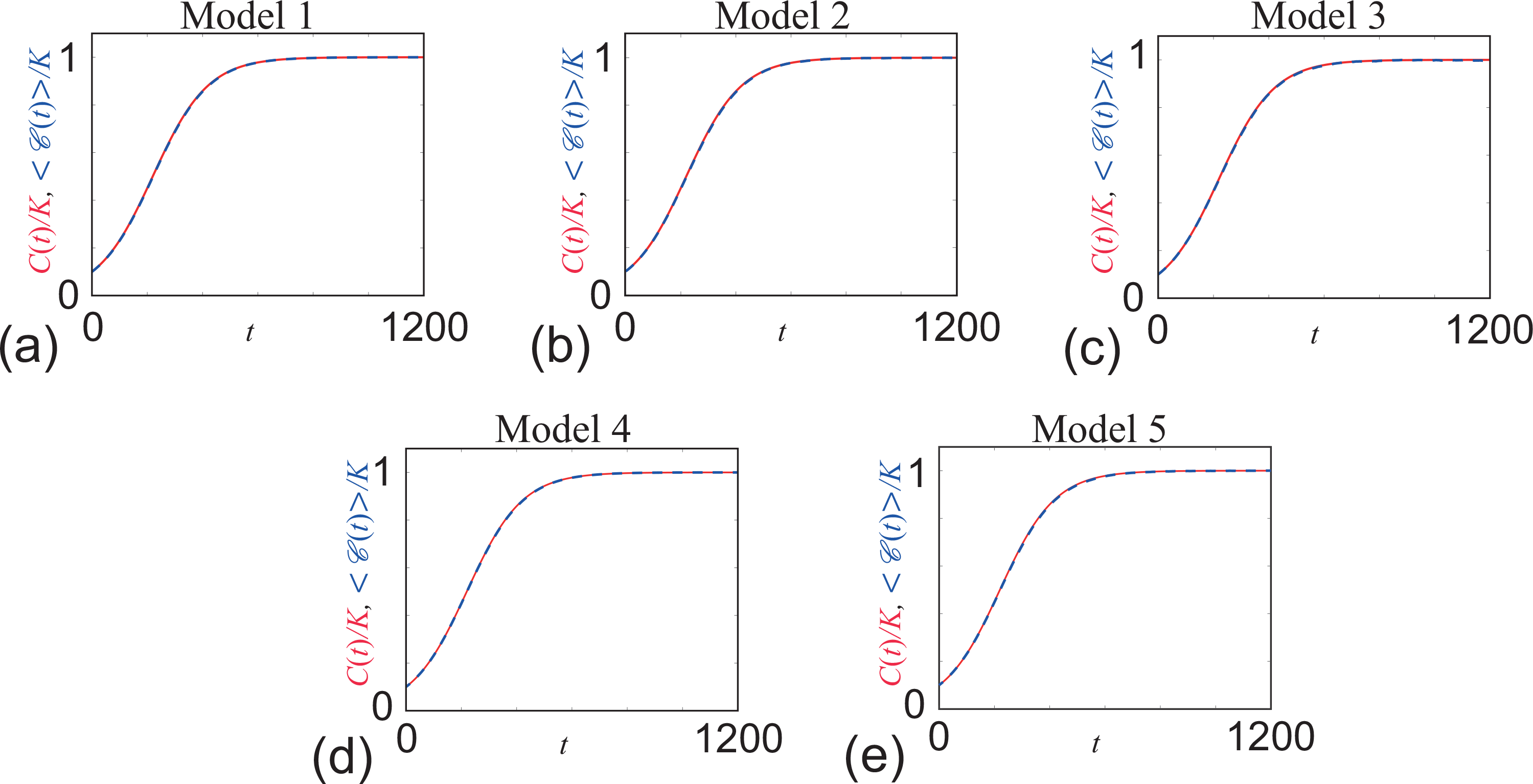
Comparison of the solution of the classical logistic equation, 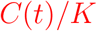, with averaged data from 1000 identically prepared realisations of the five different stochastic models, 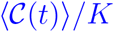. The continuum-discrete match for models 1-5 are shown in (a)-(e), respectively, with 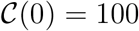, *K* = 1000 and *λ* = 0.01. /h. Parameters in the five discrete models are: *b*_1_ = 0.01 /h, *κ*_1_ = 1000; *b*_2_ = 0.0125 /h, *d*_2_ = 0.0025 /h, *κ*_2_ = 1250; *b*_3_ = 0.0125 /h, *d*_3_ = 0.0025 /h, *a*_3_ = 0.00001 agents/h; *L* = 10, Δ = 1, *b*_4_ = 0.01 /h, *m*_4_ = 1 /h; *L* = 10, Δ = 1, *b*_5_ = 0.0125 /h, *d*_5_ = 0.0025 /h, *m*_5_ = 1 /h.

Before we explore the question of whether it is possible to reliably distinguish between the five different stochastic models, it is worthwhile to point out that all plots in Figures 2 and 3 show the independent variable for 0 ≤ *t* ≤ 1200 h. While in principle it is possible to perform simulations for a longer period, the solution of Equation (1) gives *C*(1200)/*K* ≈ 0.9999 for *λ* = 0.01 /h with *C*(0)/*K* = 0.1. Therefore, this truncated time interval captures virtually the entire dynamics of the population and we do not consider any longer periods of time for this choice of *λ* and *C*(0).

Overall, the results in Figure 3 have two important consequences:

1. for each discrete model, we see that 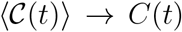 as the number of identically prepared realisations of the discrete model becomes sufficiently large, as expected (Baker and Simpson 2010). Additional results (Supplementary Material) confirms that this is also true for larger *λ*. Additional results (not shown) confirm that this also holds when we vary *C*(0);
2. if we consider generating 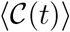 using a large number of identically prepared realisations, it is not possible to use this averaged data to distinguish which of the five very different stochastic models gave rise to that averaged population data.

### 3.2 Individual-level distinguishability

Given these observations, the task in this study is to attempt to distinguish between the five different stochastic models. The question of *model selection* is becoming increasingly important in the field of mathematical biology (Jin et al. 2017) and mathematical ecology (Johnson and Omland, 2004). For example, Sarapata and de Pillis (2014) compare a range of continuum sigmoid growth laws with data describing the temporal growth of various types of tumours and illustrate that some of the experimental datasets can be modelled using more than one choice of continuum model. Gerlee (2013) presents a thorough discussion of several different continuum models that are used to predict and explain tumour growth. Similarly, West et al. (2001) attempts to define a universal growth law to describe temporal tumour growth in various animal models. While these three previous studies focus on ordinary differential equations and temporal behaviour, similar and more complicated questions of model selection are relevant when considering combined spatial and temporal processes, such as collective cell spreading (Treloar et al. 2014). These questions have been partially explored in the context of modelling collective cell spreading processes by attempting to select the best model among a suite of distinct continuum partial differential equation models (e.g. Maini et al. 2004a; Maini et al. 2004b; Sengers et al. 2007; Sherratt and Murray 1990; Warne et al. 2018).

These previous studies in model selection are very different to the problem that we consider in this work. In these previous studies the authors compare a suite of distinct continuum models and attempt to use experimental data to select the most appropriate model. In contrast, here we consider a suite of different stochastic models, each of which has the same continuum limit description, and we attempt to distinguish between the different stochastic models. Therefore, our task is to select between different stochastic models when the continuum limit is mathematically indistinguishable. This is a challenging task, especially when we consider that standard protocols in the mathematical biology literature for interpreting results from stochastic models is to construct averaged model results by averaging results from many identically prepared realisations of the stochastic model (e.g. Baker et al. 2010; Bruna and Chapman 2012; Bruna and Chapman 2014; Dyson et al. 2013; Flegg et al. 2013; Keeling and Eames 2005). Similarly, standard protocols for interpreting noisy experimental data is to deal with averaged quantities (Jin et al. 2016; Melica et al. 2014; Pozzobon and Perré 2018; Sengers et al. 2007; Vo et al. 2015). Clearly, averaging provides no advantage when the continuum limit description of various discrete models are identical. To deal with this complication we will take the opposite approach and instead of focusing on averaging noisy data, we will examine properties of the *process noise* and explore the extent to which the inherent process noise facilitates model selection.

Here, we focus on the process noise associated with each stochastic model. To do this we consider the quantity

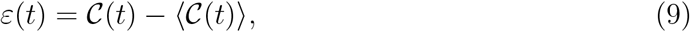

which is a time-dependent measure of the process noise that can be calculated and visualised for any realisation of any one of the five stochastic models we consider. Typical plots of *ɛ*(*t*) are given in Figure 4 for each of the five stochastic models, and in each case we show four different realisations of *ɛ*(*t*) for each model. Clearly each realisation of *ɛ*(*t*) acts like a random variable, however some model-specific trends in *ɛ*(*t*) are clear. One obvious trend is that the value of *ɛ*(*t*) for Model 3 appears to deviate further from zero than all other models. Another trend in the *ɛ*(*t*) data is that *ɛ*(*t*) for Model 1 and Model 4 appear to decay to zero at late times for each realisation whereas *ɛ*(*t*) for Models 2, 3 and 5 does not. Our aim now is to explore whether it is possible to use these individual-level differences to provide a probabilistic distinction between the five models. Furthermore, we will also explore which features of the *ɛ*(*t*) signal to use to make this distinction as clear as possible. To this end we will take a Bayesian approach (Gelman et al. 2003; Warne et al. 2019b).

### 3.3 Bayesian approach to distinguish models using process noise

As we will show, the key aspect of using process noise to distinguish between the five stochastic models is to use the process noise signal, *ɛ*(*t*), to construct summary statistics to facilitate making a reliable distinction. For the purpose of clarity we will first illustrate some key features using a simple summary statistic, and then consider refining our choice of summary statistic to refine our model selection. Perhaps the simplest way to summarise the noise signature is use the maximum deviation from zero noise,

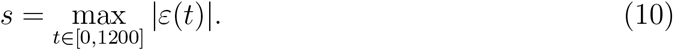

**Figure 4:**
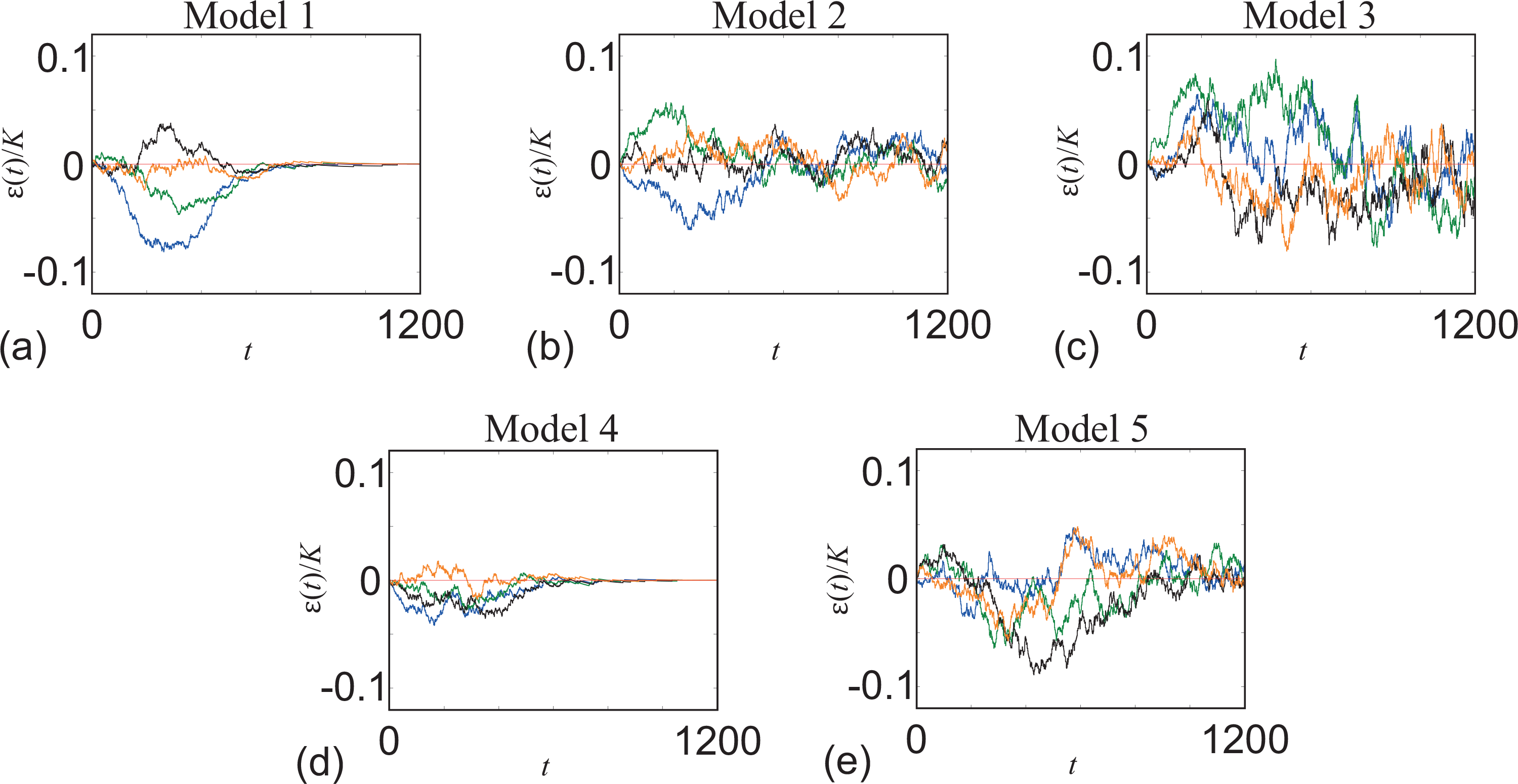
Time series data showing *ɛ*(*t*)/*K* for four realisations for each of the five models. The time series data for models 1-5 are shown in (a)-(e), respectively, with 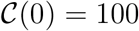, *K* = 1000 and *λ* = 0.01. For each stochastic model, four typical realisations of *ɛ*(*t*) are shown in green, blue, black and orange.

This choice of summary statistic allows us to replace each time series, *ɛ*(*t*), with a single scalar value, *s*, and we will now explore the extent to which using this information enables us to reliably distinguish between the five stochastic models. To achieve this we take the 1000 identically prepared realisations of each stochastic model that we used to construct averaged density data in Figure 3 and we calculate *s* for each of the realisations. Using MATLABs `ksdensity` function (Mathworks, 2019b) we convert the discrete distribution of *s* for each model into a smooth, approximate density distribution, as shown in Figure 5.

**Figure 5:**
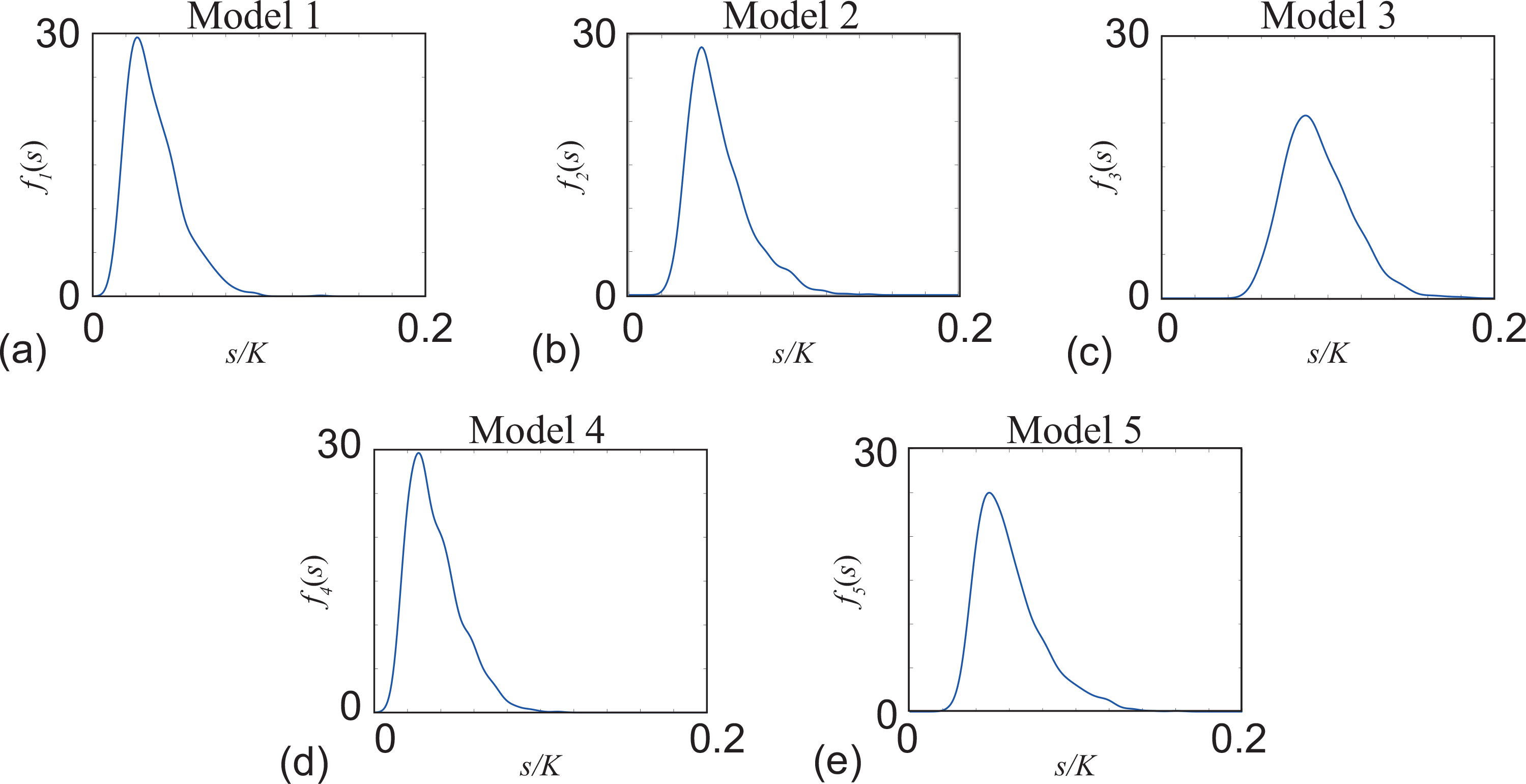
Univariate density profiles for the five stochastic models with the simple univariate summary statistic, Equation (10). (a)-(e) show density estimates, *f*_*i*_(*s*), as a function of the univariate summary statistic, *s*, given by Equation (10). Each density profile is constructed using 1000 identically prepared realisations as describing in Figure 3, and the smoothed density profile is obtained with MATLABs *ksdensity* function (Mathworks, 2019b). All results correspond to *λ* = 0.01 /h.

A qualitative comparison of the density profiles in Figure 5 confirms that these density profiles capture some of the putative differences that we described in Figure 4. For example, if we consider a maximum deviation of *s/K* = 0.1, the density associated with Model 1 is relatively small, *f*_1_(0.1) ≈ 0, whereas the density for Model 3 is relatively large, *f*_3_(0.1) ≈ 20. To make use of this difference in a Bayesian framework, suppose that we are given *f*_*i*_(*s*) for *i* = 1, 2, …, 5. If we then perform a single realisation of a randomly-selected model and calculate *s* from that realisation we apply Bayes theorem to give

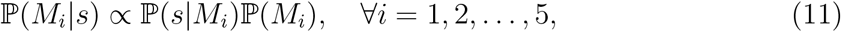

where ℙ(*M*_*i*_|*s*) is the probability of model *i* given the summary statistic *s*, ℙ(*s*|*M*_*i*_) is the probability density of summary statistic *s* given model *i*, and ℙ(*M*_*i*_) is the prior model probability, which encodes our previous knowledge of which model is most appropriate for the data.

For all of our work we make the conservative assumption that the prior specifies that all models are equally likely, ℙ(*M*_*i*_) = 1/5 for *i* = 1, 2, …, 5. We note that ℙ(*s*|*M*_*i*_) is the likelihood, and in this case we have ℙ(*s*|*M*_*i*_) = *f*_*i*_(*s*) for *i* = 1, 2, …, 5. This means that for a single realisation of one of the five models chosen at random ℙ(*M*_*i*_|*s*) is simply proportional to *f*_*i*_(*s*), and we can calculate ℙ(*M*_*i*_|*s*) for each *i* = 1, 2, …, 5 by ensuring that 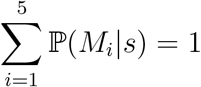. Again, for all our work, we construct the likelihood using a summary statistic, *s*, instead of the data, *ɛ*(*t*). This choice is made on practical grounds as it is computationally efficient to work with lower dimensional summary statistics to summarise the data than it is to work with all data collected to construct *ɛ*(*t*) (Maclaren et al. 2015; Maclaren et al. 2017; Lambert et al. 2018).

Instead of considering just one single realisation of a particular model chosen at random, a more realistic scenario is that we consider a small number of identically prepared realisations of a particular model. Since each identically prepared realisation of the unknown model, and the associated summary statistic, are independent, we obtain

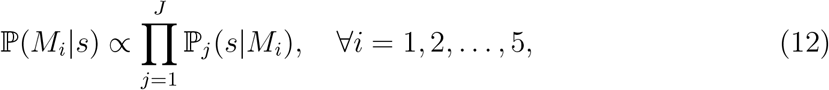

under the conservative assumption that the prior specifies that each model is equally likely. Here, ℙ_*j*_(*s*|*M*_*i*_) is the probability density that the summary statistic *s*, in the *j*^th^ identically prepared realisation is associated with model *i*. In this formulation we are considering *J* identically prepared realisations. Again, with this information we can calculate ℙ(*M*_*i*_|*s*) for each *i* = 1, 2, …, 5 by ensuring that 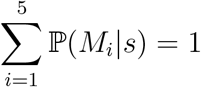.

To demonstrate the performance of this simple summary statistic we will now consider each of the five stochastic models in turn. For each model, *i* = 1, 2, …, 5, we take *J* = 5 identically prepared realisations of a randomly-chosen model to calculate ℙ(*M*_*i*_|*s*) according to Equation (12). We repeat this process 1000 times, giving us access to a distribution of estimates for ℙ(*M*_*i*_|*s*), from which we can calculate the sample mean and the 95% credible interval. Results in Figure 6(a) show ℙ(*M*_*i*_|*s*), for *i* = 1, 2, …, 5, for Model 1. Here, point estimates correspond to the sample mean of ℙ(*M*_*i*_|*s*) and the error bars denote the 95% credible interval. Figure 6(a) includes a horizontal line at 1/5, indicating the prior distribution. In this case we see that ℙ(*M*_2_*|s*), ℙ(*M*_3_|*s*) and ℙ(*M*_5_|*s*) are all smaller than 1/5, whereas ℙ(*M*_1_|*s*) and ℙ(*M*_4_|*s*) are greater than the prior values. This means that the simple summary statistic correctly indicates that Models 2, 3 and 5 are less likely to explain the summary statistic than Models 1 and 4. However, this process gives ℙ(*M*_1_|*s*) ≈ ℙ(*M*_4_|*s*), indicating that this summary statistic does not reliably distinguish between Models 1 and 4.

**Figure 6:**
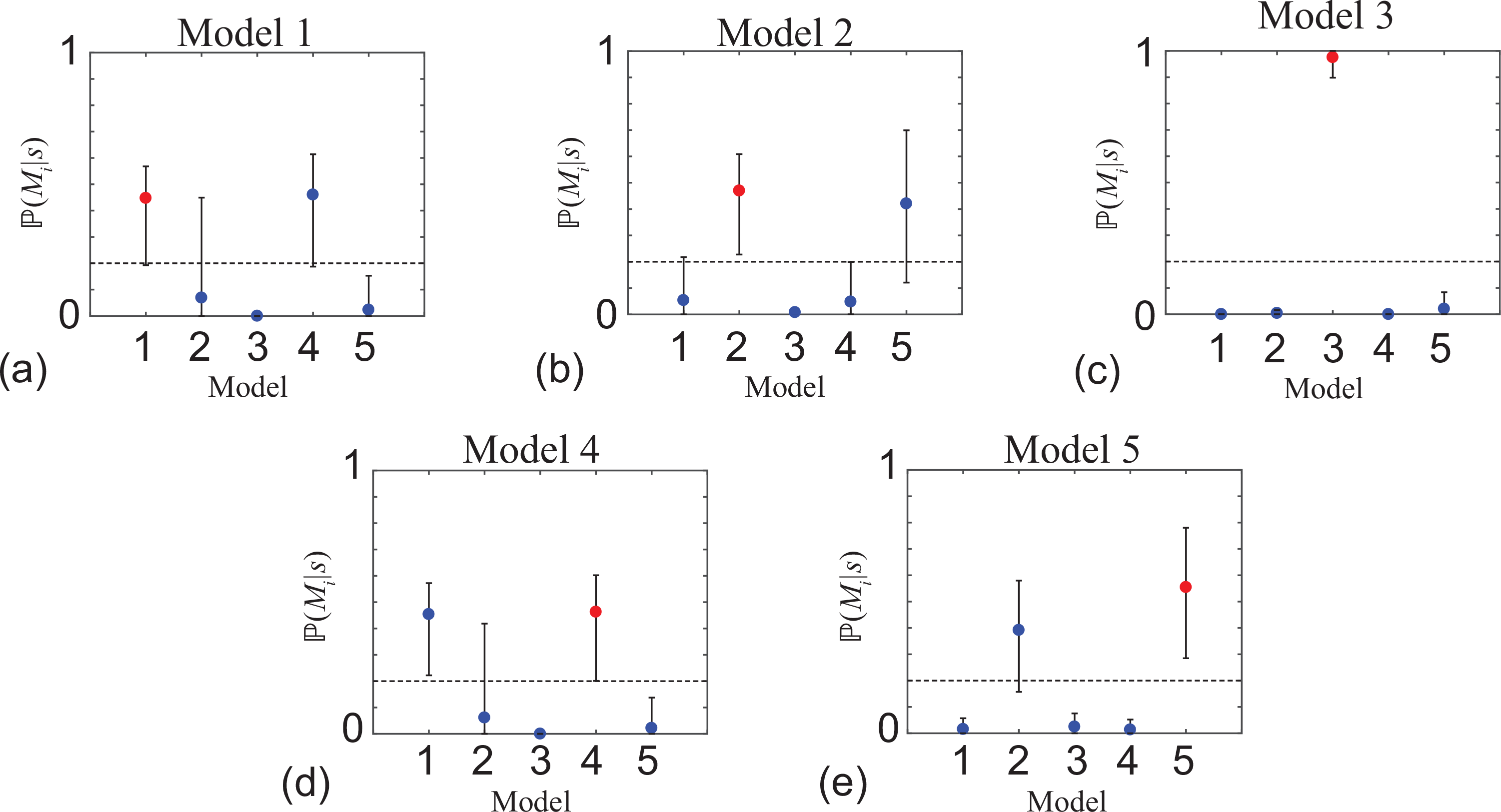
Model identification with the simple univariate summary statistic, *s*, given by Equation (10). Results in (a)-(e) show point estimates of ℙ(*M*_*i*_|*s*) for *i* = 1, 2, …, 5 and the uncertainty in the estimate is indicated by the error bars. The point estimates correspond to the sample mean and the error bar corresponds to the sample mean plus or minus one sample standard deviation, both calculated using 1000 identically prepared estimates of ℙ(*M*_*i*_|*s*) for *i* = 1, 2, …, 5 with *J* = 5. Each subfigure shows a horizontal line at 1/5, indicating the prior distribution of ℙ(*M*_*i*_) for *i* = 1, 2, …, 5. In each subfigure, maximum value of 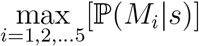 is shown in red, and in each case this is the correct model choice. All results correspond to *λ* = 0.01 /h.

This process of calculating ℙ(*M*_*i*_|*s*), for *i* = 1, 2, …, 5 for each of the candidate models is repeated in Figure 6(b)-(e) for Models 2, 3, 4 and 5, respectively. Results in Figure 6(b) explore the ability of the simple summary statistic to distinguish Model 2 and we see that the process correctly identifies the Models 1, 3 and 5 are less likely to explain the data whereas Models 2 and 5 both give similar results. Results in Figure 5(c) explore the ability of the simple summary statistics to identify Model 3, and in this case we have a very promising result that ℙ(*M*_3_|*s*) ≈ 1, and all other candidate models have close to zero probability. Results in Figure 6(d) and (e) are similar to the results in Figure 6(a) and (b) since the simple summary statistic correctly assigns a lower probability to three of the candidate models, but is unable to reliably distinguish between two other candidate models.

In addition to simply calculating the sample mean and 95% credible intervals of our estimates of ℙ_*j*_(*M*_*i*_|*s*), as reported in Figure 6, we can also visualise the entire distribution of 1000 estimates of ℙ_*j*_(*M*_*i*_|*s*). Results in Figure 7 show histograms of ℙ_*j*_(*M*_*i*_|*s*) for Models 1, 2, …, 5, respectively. In each case, we see that each histogram appears to be unimodal, which indicates that summarising these distributions in terms of the sample mean is useful and we see that some of the distributions, such as the histogram in Figure 7(c), are more peaked than the others, indicating a higher certainty in these results.

**Figure 7:**
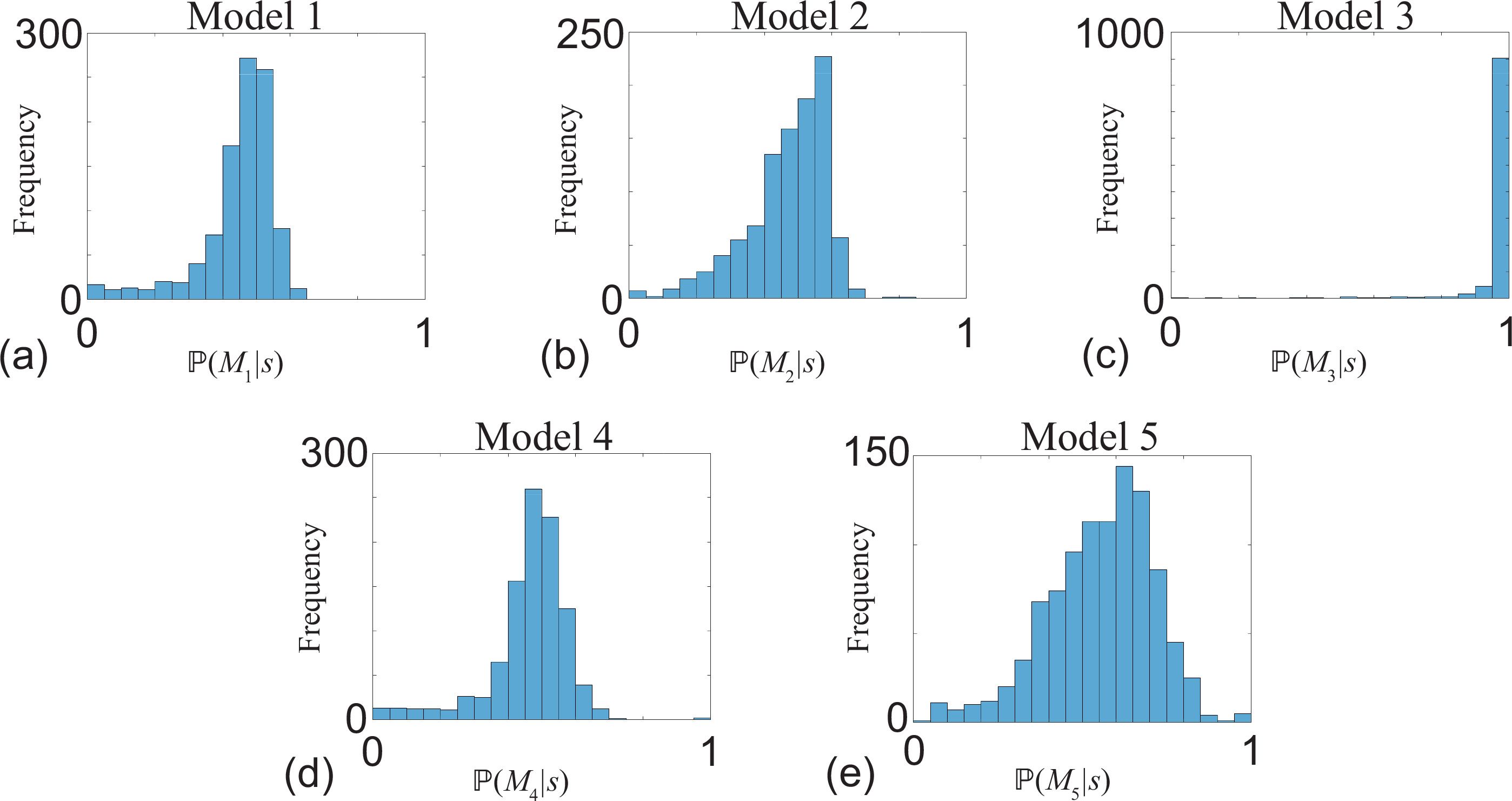
Distribution of ℙ(*M*_*i*_|*s*) using the simple univariate summary statistic, *s*, given by Equation (10). Results in (a)-(e) show distributions of estimates of ℙ(*M*_*i*_|*s*) calculated using 1000 identically prepared estimates of ℙ(*M*_*i*_|*s*) for *i* = 1, 2, …, 5 with *J* = 5. All results correspond to *λ* = 0.01 /h.

Results in Figures 6-7 are promising. The simplest possible summary statistic, Equation (10), can partially distinguish between the five stochastic models that are completely indistinguishable when we consider averaged data in Figures 2-3. We now explore our ability to improve these preliminary results by refining the summary statistic. A key feature of *ɛ*(*t*), clearly evident in Figure 4, is that the temporal noise signature appears to depend on time. This suggests that improved results might be obtained by constructing a more detailed summary statistic that characterises both *early* and *late* features of *ɛ*(*t*). To this end we partition the time interval considered, 0 < *t* < 1200, into quartiles, and collect data describing the noise signature in the first quartile, 0 < *t* < 300, and the fourth quartile, 900 < *t* < 1200. Again, we summarise these two process noise signatures by the maximum deviation from zero, giving 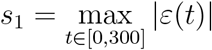 and 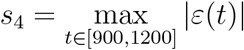. This means that instead of summarising the noise signature using a single number, *s*, we now summarise each stochastic simulation using vector with two components, (*s*_1_, *s*_4_). Using the 1000 identically prepared realisations of each model to construct averaged density data in Figure 3 we calculate (*s*_1_, *s*_4_) for each and use the `ksdensity` function in MATLAB (Mathworks, 2019b) to form smooth, approximate bivariate density distributions associated with each stochastic model, as shown in Figure 8. Results in Figure 8 confirm that we have visually obvious differences in the bivariate distribution of (*s*_1_, *s*_4_). Most notably these distributions clearly show the larger fluctuations inherent in Model 3, and we see that the late process noise in Models 1 and 4 are very different to the late process noise in Models 2 and 5.

**Figure 8:**
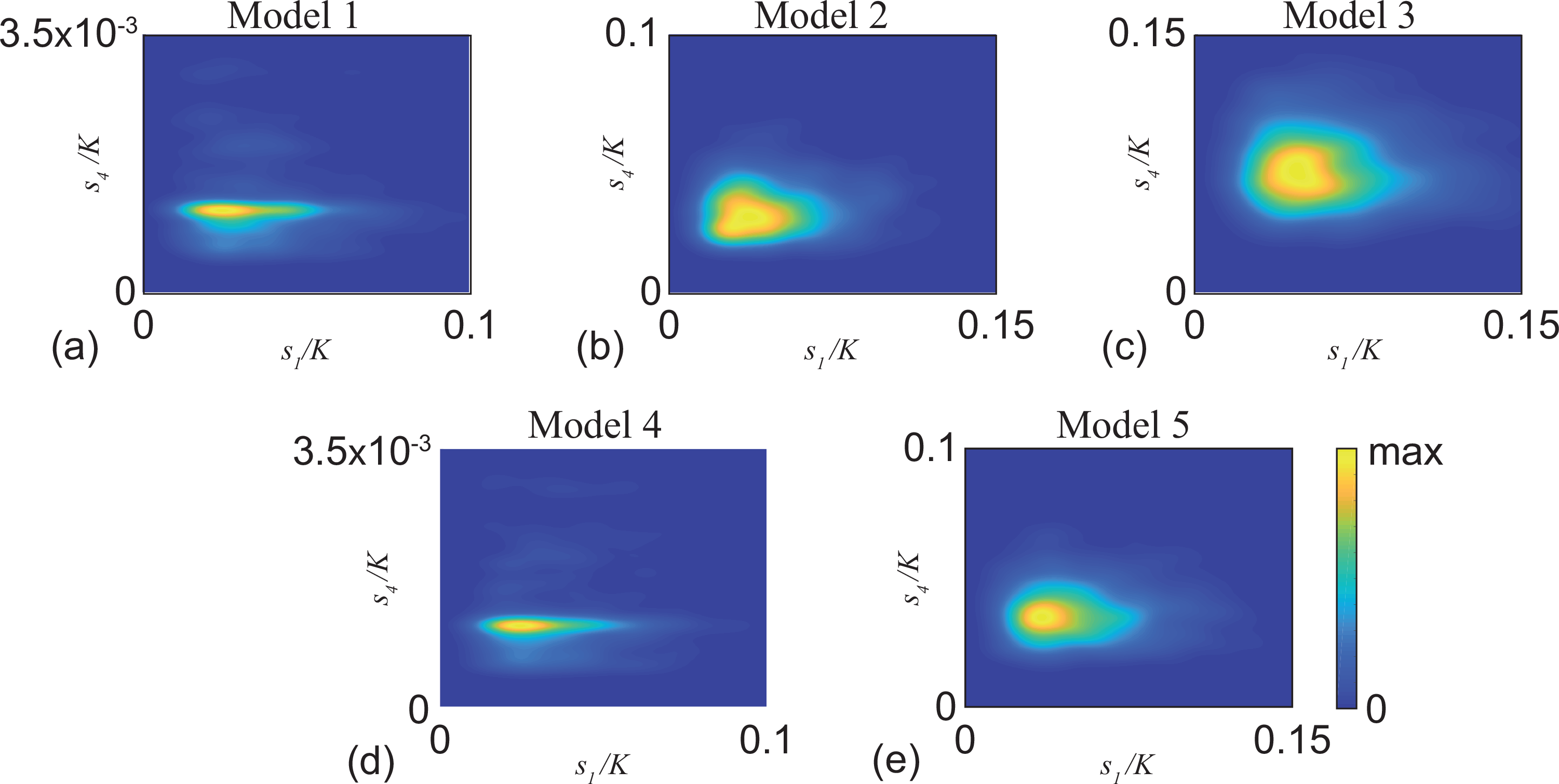
Bivariate density profiles for the five stochastic models with 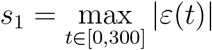 and 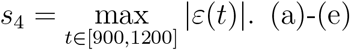. show density estimates, *f*_*i*_(*s*_1_, *s*_4_), shown as a function of *s*_1_/*K* and *s*_4_/*K*, for *i* = 1, 2, …, 5, respectively. Each density profile is constructed using 1000 identically prepared realisations as describing in Figure 3, and the smoothed bivariate density profile is obtained with MATLABs ksdensity function (Mathworks, 2019b). All results correspond to *λ* = 0.01 /h.

Using the refined summary statistic densities in Figure 8, we estimate ℙ(*M*_*i*_|*s*) for each model by taking *J* = 5 identically prepared realisations of a randomly-chosen model according to Equation (12). Again, we repeat this process 1000 times, giving us a distribution of estimates for ℙ(*M*_*i*_|*s*), from which we can calculate the sample mean and the 95% credible interval. Results in Figure 9(a) show ℙ(*M*_*i*_|*s*), for *i* = 1, 2, …, 5, for Model 1. As for the results based on the simple summary statistic in Figure 6(a), the refined summary statistic gives ℙ(*M*_2_|*s*), ℙ(*M*_3_|*s*) and ℙ(*M*_5_|*s*) are all smaller than 1/5, whereas ℙ(*M*_1_|*s*) and ℙ(*M*_4_|*s*) are greater than the prior values. This result correctly implies that the bivariate summary statistic assigns a low probability to Models 2, 3 and 5, and a higher probability to Models 1 and 4.

**Figure 9:**
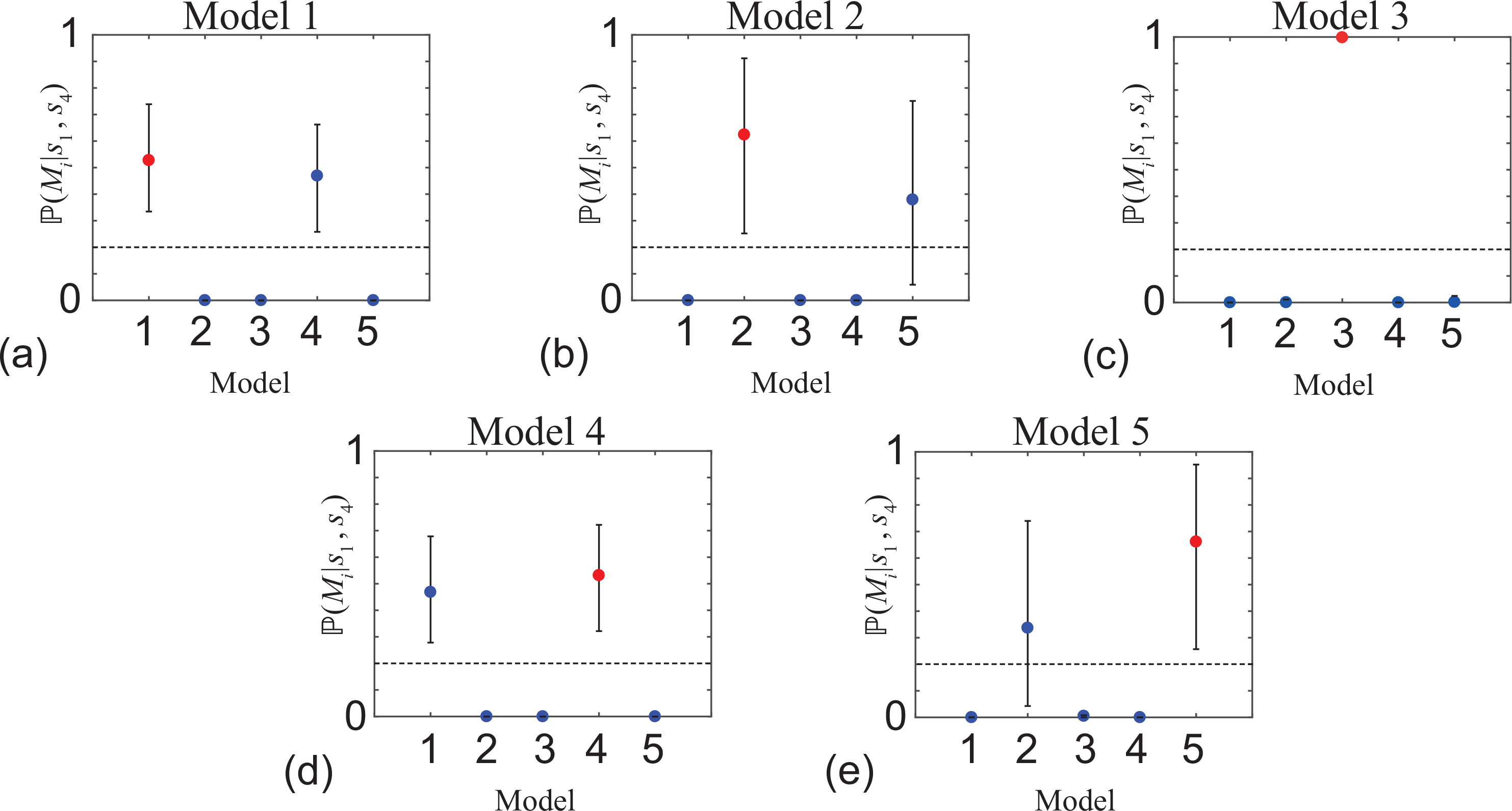
Model identification with the bivariate summary statistic, (*s*_1_, *s*_2_). Results in (a)-(e) show point estimates of ℙ(*M*_*i*_|*s*) for *i* = 1, 2, …, 5 and the uncertainty in the estimate is indicated by the error bars. The point estimates correspond to the sample mean and the error bar corresponds to the sample mean plus or minus one sample standard deviation, both calculated using 1000 identically prepared estimates of ℙ(*M*_*i*_|*s*) for *i* = 1, 2, …, 5 with *J* = 5. Each subfigure shows a horizontal line at 1/5, indicating the prior distribution of ℙ(*M*_*i*_) for *i* = 1, 2, …, 5. In each subfigure, maximum value of 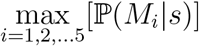 is shown in red, and in each case this is the correct model choice. All results correspond to *λ* = 0.01 /h.

Comparing results in Figure 6 and Figure 9 confirms that the additional information in the bivariate summary statistic leads to an improved ability to distinguish between some of the models. For example, if we compare results in Figure 6(b) with the results in Figure 9(b) we see that both summary statistics give point estimates of ℙ(*M*_*i*_|*s*) ≈ 0 and ℙ(*M*_*i*_|*s*_1_, *s*_4_) ≈ 0 for *i* = 1, 3 and 4, correctly identifying that Models 1, 3 and 4 are unlikely to explain the data. Results in Figure 6(b) indicate that the simple univariate summary statistic leads to point estimates with ℙ(*M*_2_|*s*) ≈ ℙ(*M*_5_|*s*) confirming that the univariate summary statistic does not reliably distinguish between Model 2 and Model 5. In contrast, results in Figure 9(b) indicate that ℙ(*M*_2_|*s*_1_, *s*_4_) > ℙ(*M*_5_|*s*_1_, *s*_4_) confirming that the bivariate summary statistic correctly indicates that Model 2 gives the best explanation of the summary statistic. Again, instead of relying simply on point estimates and quartile distributions in Figure 9 for the improved summary statistic, results in Figure 10 show the distributions of ℙ(*M*_*i*_|*s*_1_, *s*_4_) for each model.

**Figure 10:**
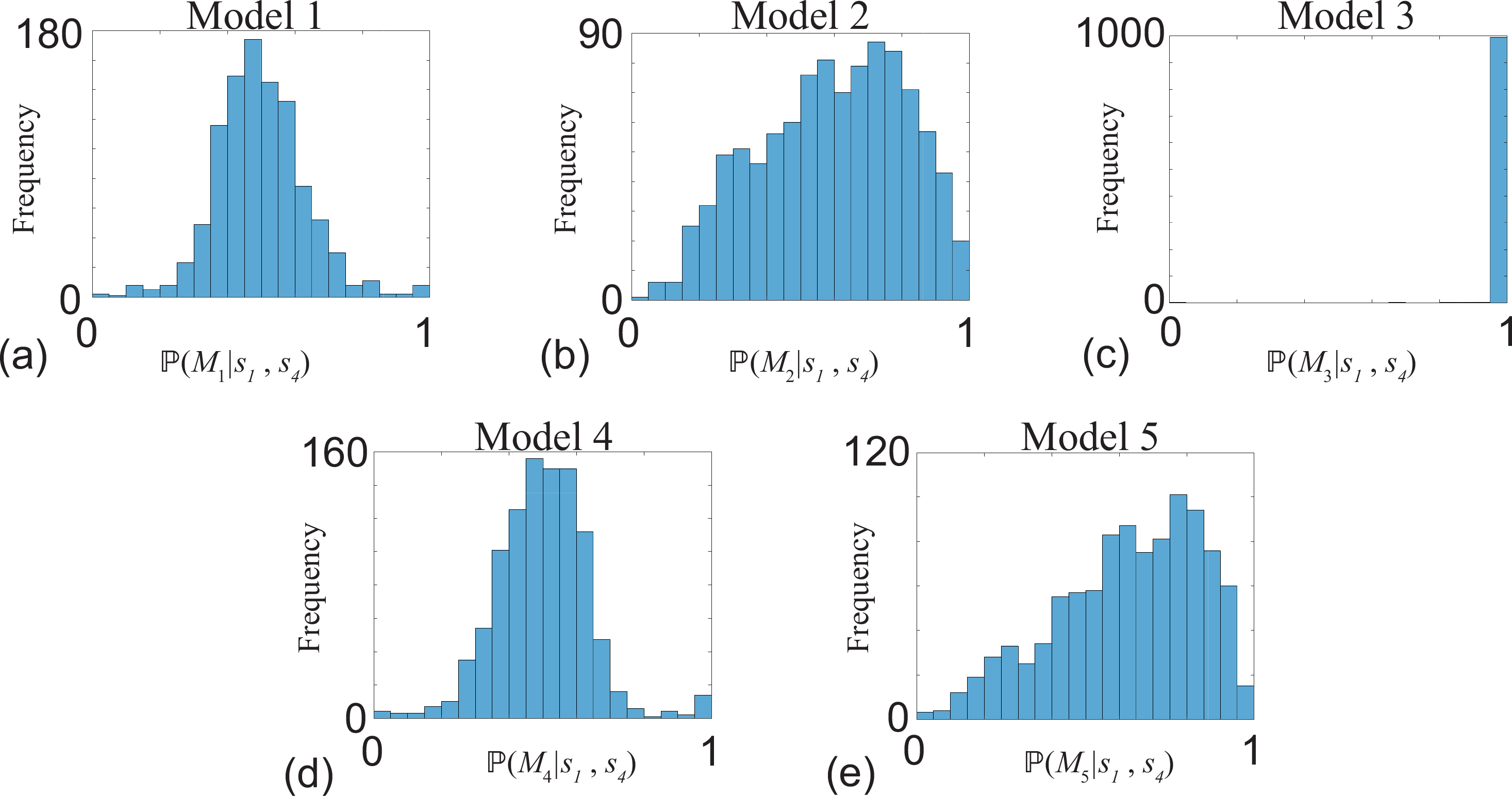
Distribution of ℙ(*M*_*i*_|*s*) using the bivariate summary statistic, (*s*_1_, *s*_4_). Results in (a)-(e) show distributions of estimates of ℙ(*M*_*i*_|*s*) calculated using 1000 identically prepared estimates of ℙ(*M*_*i*_|*s*) for *i* = 1, 2, …, 5 with *J* = 5. All results correspond to *λ* = 0.01/h.

## Conclusions

The process of model selection is becoming increasingly of interest to the mathematical biology community. Unlike other areas of applied mathematics, such as fluid mechanics and solid mechanics, where there is consensus about what kinds of models are most appropriate, in mathematical biology it is often the case that several competing models are available to describe similar phenomena (Gerlee 2014; Sarapata and de Pillis 2014; Jin et al. 2016; Browning et al. 2017). The process of model selection often involves taking a set of observations and exploring the extent to which results from a suite of continuum models provide the best explanation of those data. Various approaches to model selection can be taken such as using maximum likelihood (Jin et al. 2014) or Bayesian approaches (Warne et al. 2018). In this work we explore a more challenging problem in model selection by working with a suite of stochastic models describing population growth processes. The main challenge is that we consider five distinct discrete models and show, through simulation, that each model is well described by the same continuum logistic growth model, given by Equations (1)-(2). If we take the usual approach and average data from many identically prepared realisations from each stochastic model that the averaged results are completely indistinguishable. This means that the usual tools used in model selection for continuum models cannot be used to distinguish between the five stochastic models that we consider.

To make progress we take a different approach and examine the *process noise* associated with each model and find that there are certain features of the process noise that appear to be different between the five different stochastic models. We demonstrate how to make a probabilistic distinction between the five models in a Bayesian framework by constructing appropriate summary statistics from the process noise. Our results show that even using the simplest possible summary statistic, the maximum deviation away from zero noise, allows us to reliably distinguish between some of the models. In particular, we find that Model 3 involving spatially implicit birth-death-annihilation is remarkably easy to distinguish from the other four models using the simplest possible summary statistic. Furthermore, we find that certain groups of models are also easy to distinguish using this simple approach. As expected, when we extend the summary statistic to include information about both the *early* and *late* portions of the process noise, we find that our ability to distinguish between the five models is enhanced. Therefore, while all five models are completely indistinguishable using population-level information, we show that individual-level information can be used in a Bayesian framework to provide a probabilistic distinction between the various models.

There are many ways in which the work we present here can be extended. The most obvious line of extension would be to consider using different summary statistics in an attempt to further improve our model selection results. While all results in Figure 6 and Figure 9 are based on taking the maximum absolute value of *ɛ*, we also explored with other choices of summary statistic, such as working with the maximum value of *ɛ*, the minimum value of *ɛ* and measures of autocorrelation in *ɛ*. These additional results (not shown) did not improve our ability to distinguish between the give models over the results in Figure 6 and Figure 9 and so we choose to present results based on the simplest summary statistic only. Another way to potentially improve our ability to distinguish between the five models is to use higher dimensional summary statistics. Just as the results in Figure 9 for the bivariate summary statistic, (*s*_1_, *s*_4_), improve our ability to distinguish between the models than the results in Figure 6 for the univariate summary statistic, *s*, we expect that introducing additional information into the summary statistic definition, and hence working with higher dimensional summary statistics, will improve our model selection results. Since we have already partially demonstrated that more detailed summary statistics can lead to improved model section in Figure 6 and Figure 9, we do not pursue further refinements along these lines here. Another point for consideration would be to relax our conservative assumption of working with uniform prior distributions and incorporating some uncertainty via the prior. A slightly different extension would be to make use of our observation that the process noise, *ɛ*(*t*), is time dependent. This suggests that it would be possible to explore the question of the optimal time(s) to make observations (Dehideniya et al. 2018; Johnston et al. 2016; Overstall and McGree 2019; Warne et al. 2017). One final avenue for further exploration would be to examine how the concepts and techniques developed here for the classical logistic growth model would apply to other stochastic models in the mathematical biology and mathematical ecology literature, such as stochastic analogues of the well known SIR and SIS disease transmission models [1]. We leave such analysis for future consideration.

