## Supplementary Material for "Process noise distinguishes between indistinguishable population dynamics"

### Supplementary Material: Process noise distinguishes between indistinguishable population dynamics

### 1 Additional Results

As we describe in the main document, all results and discussion in the main document correspond to logistic growth with $\lambda = 0.01$ /h, which is a lower bound on the growth rate of biological cells. Here we provide additional results, using the same format to present the results and the same nomenclature for $\lambda = 0.05$ /h. This higher growth rate is an upper bound for the growth rate of biological cells. The main difference between the results in the main document and the supplementary material document is the growth rate, $\lambda$. This change means that we also alter the time duration in the stochastic simulations. We note that the solution of Equation (1) (main document) gives $C(240)/K \approx 0.9999$ for $\lambda = 0.05$ /h with $C(0)/K = 0.1$. Therefore, since we increase proliferation rate by a factor of five, we reduce the duration of the simulations by a factor of five since virtually all the population dynamics occur in the interval $0 \leq t \leq 240$ h for the higher proliferation rate. This means that when we construct the first univariate summary statistic from the stochastic model we define $s = \max_{t \in [0,240]} |\varepsilon(t)|$, and when we take the second bivariate summary statistic we define $s_1 = \max_{t \in [0,60]} |\varepsilon(t)|$ and $s_4 = \max_{t \in [180,240]} |\varepsilon(t)|$.

We now demonstrate that averaged data from the five stochastic models matches Equations (1)-(2) (main document). Results in Figure 1 are presented in the exact same format as in Figure 3 (main document) except here we consider $\lambda = 0.05$ /h. These additional results show that averaged data in Figure 1 are indistinguishable from the solution of Equation (1) (main document).

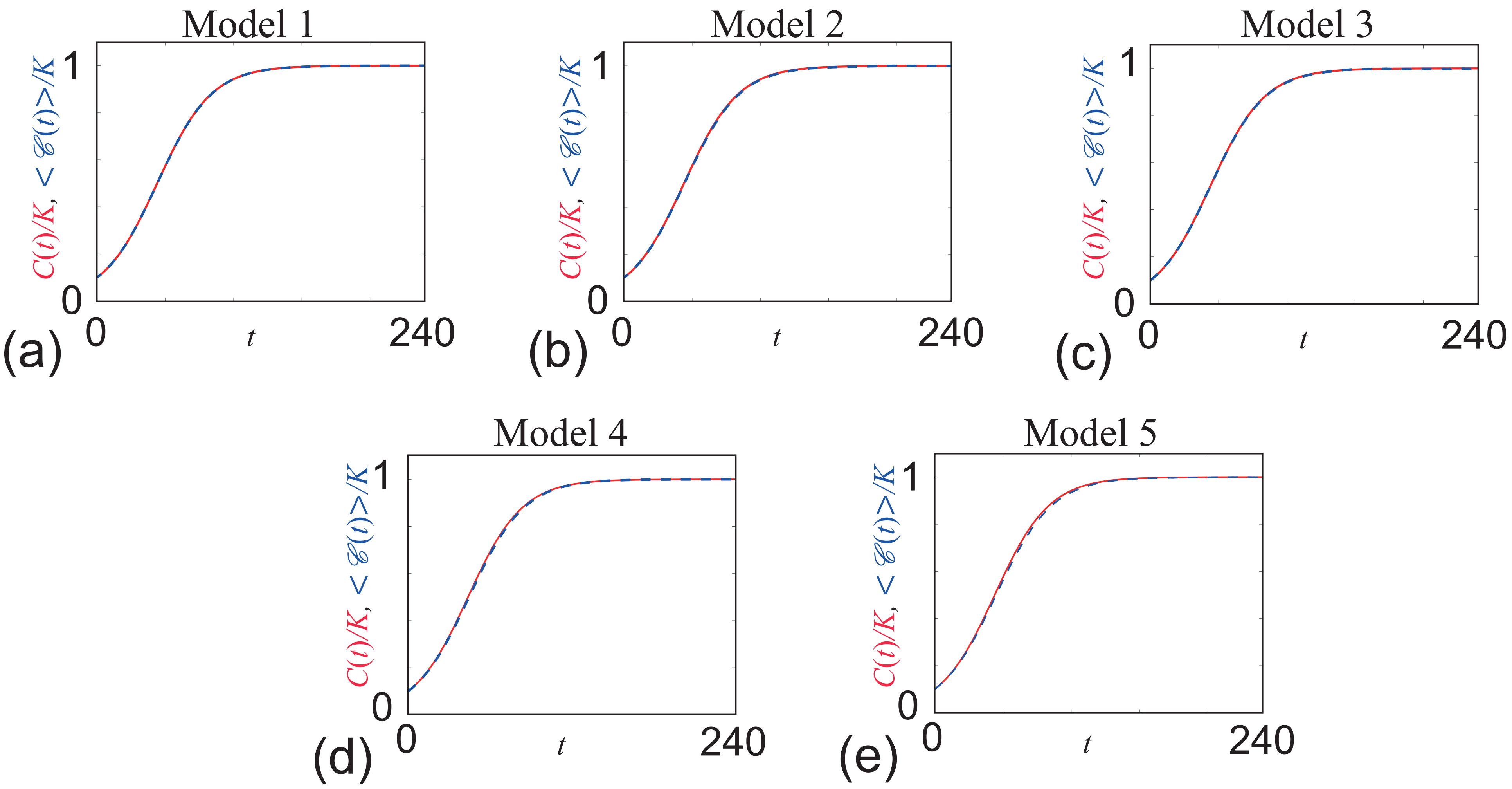


Figure 1: **Comparison of the solution of the classical logistic equation, $C(t)/K$, with averaged data from 1000 identically prepared realisations of the five different stochastic models, $\langle \mathcal{C}(t) \rangle / K$.** The continuum-discrete match for models 1-5 are shown in (a)-(e), respectively, with $\mathcal{C}(0) = 100$, $K = 1000$ and $\lambda = 0.05$. /h. Parameters in the five discrete models are: $b_1 = 0.05$ /h, $\kappa_1 = 1000$; $b_2 = 0.0625$ /h, $d_2 = 0.0125$ /h, $\kappa_2 = 1250$; $b_3 = 0.0625$ /h, $d_3 = 0.0125$ /h, $a_3 = 0.00001$ agents/h; $L = 10$, $\Delta = 1$, $b_4 = 0.05$ /h, $m_4 = 1$ /h; $L = 10$, $\Delta = 1$, $b_5 = 0.0625$ /h, $d_5 = 0.0125$ /h, $m_5 = 1$ /h.

Given that averaged data from the five stochastic models are indistinguishable from the solution of Equation (1) (main document), we now explore the properties of the process noise for the higher proliferation rate. Results in Figure 2 show four plots of $\varepsilon(t)$ (Equation (9) main document) for each of the five stochastic models we consider. Similar to the results in the main document, here we see some trends in the $\varepsilon(t)$ data that could be used to distinguish between the five models. One obvious trend in the $\varepsilon(t)$ data is that the value of $\varepsilon(t)$ for Model 3 appears to deviate further from $\varepsilon(t) = 0$ than all other models. Another trend in the $\varepsilon(t)$ data is that $\varepsilon(t)$ for Model 1 and Model 4 appear to decay to zero at late time for each realisation whereas the $\varepsilon(t)$ for Models 2, 3 and 5 do not.

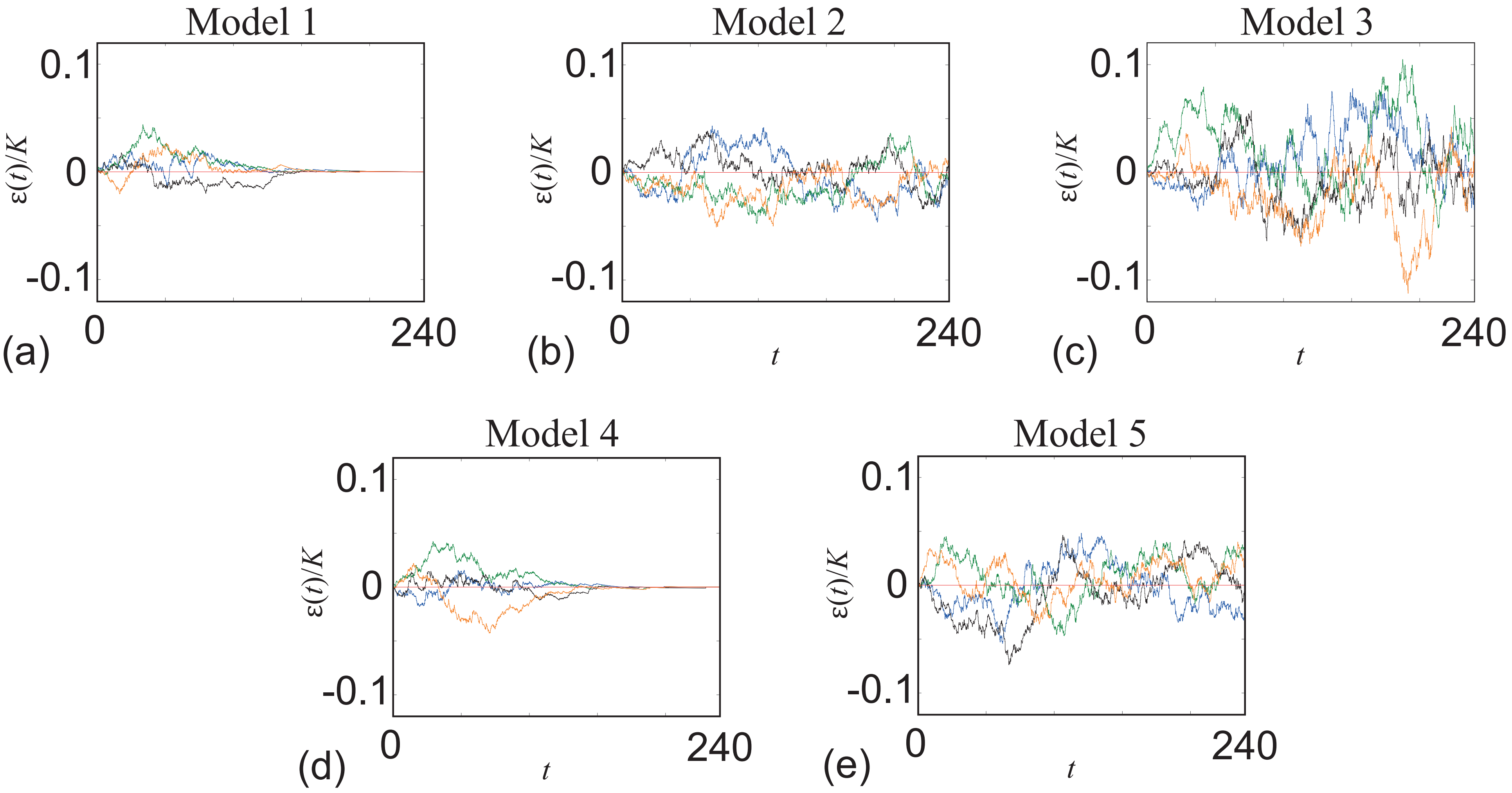


Figure 2: **Time series data showing $\varepsilon(t)/K$ for four realisations for each of the five models.** The time series data for models 1-5 are shown in (a)-(e), respectively, with $\mathcal{C}(0) = 100$, $K = 1000$ and $\lambda = 0.05$ /h. For each stochastic model, four typical realisations of $\varepsilon(t)$ are shown in green, blue, black and orange.

With this $\varepsilon(t)$ data in Figure 2 we use MATLABs `ksdentiy` routine to form five univariate distribution functions, $f_i(s)$ for $i = 1, 2, \ldots, 5$ (not shown). We then use the same procedure as outlined in the main document to calculate $\mathbb{P}(M_i|s)$ for $i = 1, 2, \ldots, 5$. Results in Figure 3 are directly analogous to the results in Figure 6 (main document).

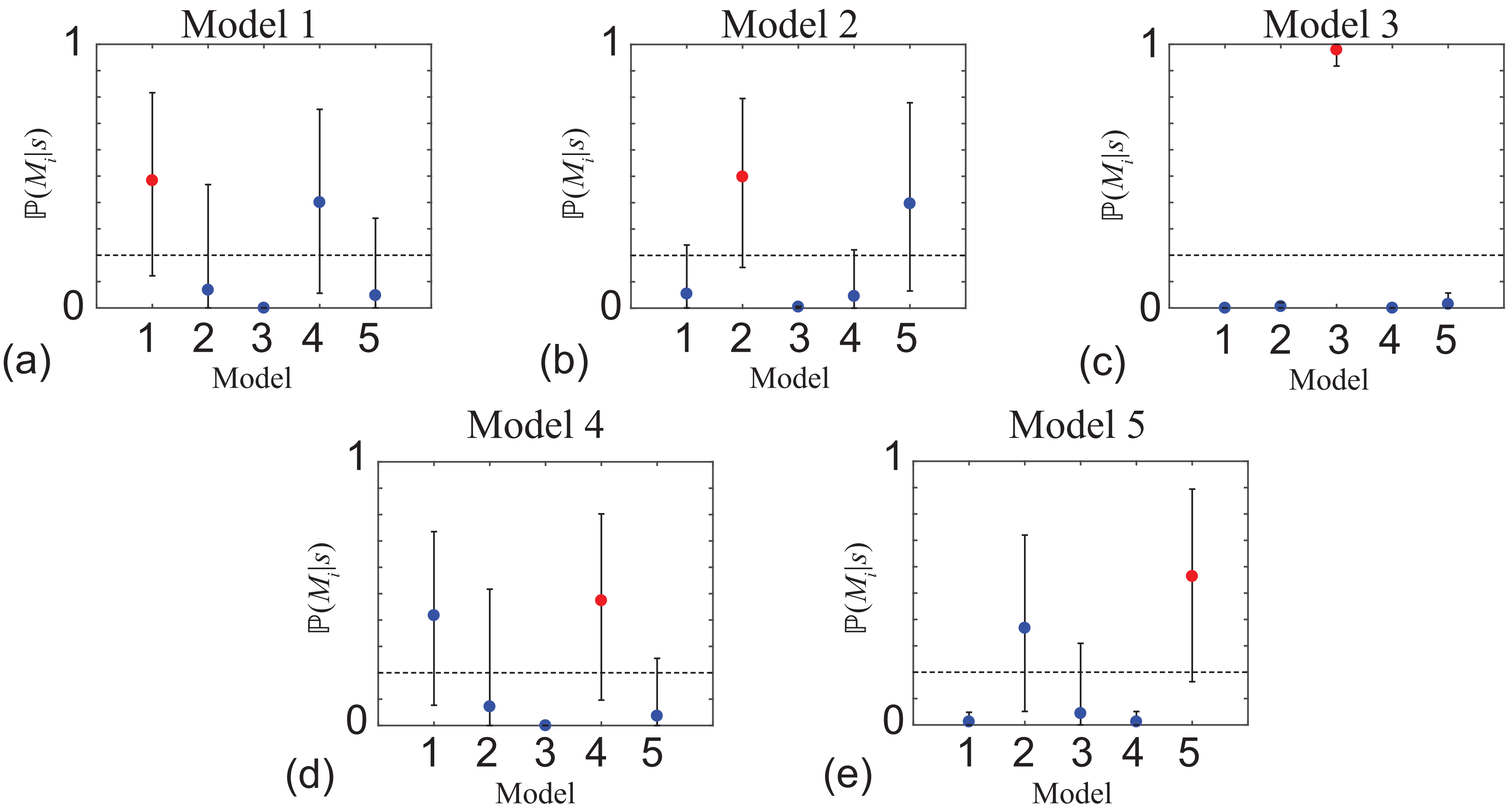


Figure 3: **Model identification with the simple univariate summary statistic, $s$, given by Equation (10) (main document).** Results in (a)-(e) show point estimates of $\mathbb{P}(M_i|s)$ for $i = 1, 2, \ldots, 5$ and the uncertainty in the estimate is indicated by the error bars. The point estimates correspond to the sample mean and the error bar corresponds to the sample mean plus or minus one sample standard deviation, both calculated using 1000 identically prepared estimates of $\mathbb{P}(M_i|s)$ for $i = 1, 2, \ldots, 5$ with $J = 5$. Each subfigure shows a horizontal line at $1/5$, indicating the prior distribution of $\mathbb{P}(M_i)$ for $i = 1, 2, \ldots, 5$. In each subfigure, maximum value of $\max_{i=1,2,\ldots 5}[\mathbb{P}(M_i|s)]$ is shown in red, and in each case this is the correct model choice. All results correspond to $\lambda = 0.05$ /h.

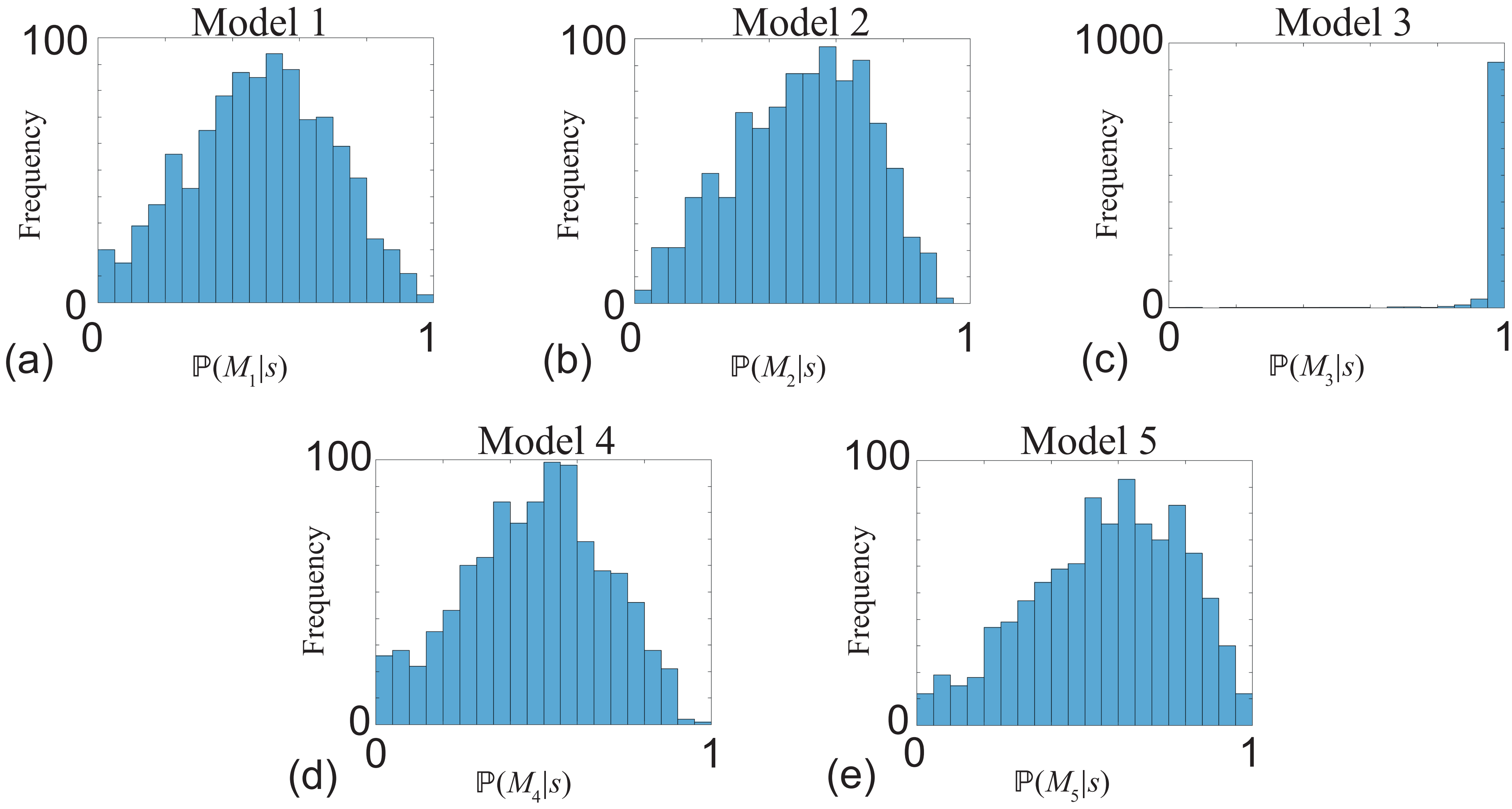


Figure 4: **Distribution of $\mathbb{P}(M_i|s)$ using the simple univariate summary statistic, $s$, given by Equation (10) (main document).** Results in (a)-(e) show distributions of estimates of $\mathbb{P}(M_i|s)$ calculated using 1000 identically prepared estimates of $\mathbb{P}(M_i|s)$ for $i = 1, 2, \ldots, 5$ with $J = 5$. All results correspond to $\lambda = 0.05$ /h.

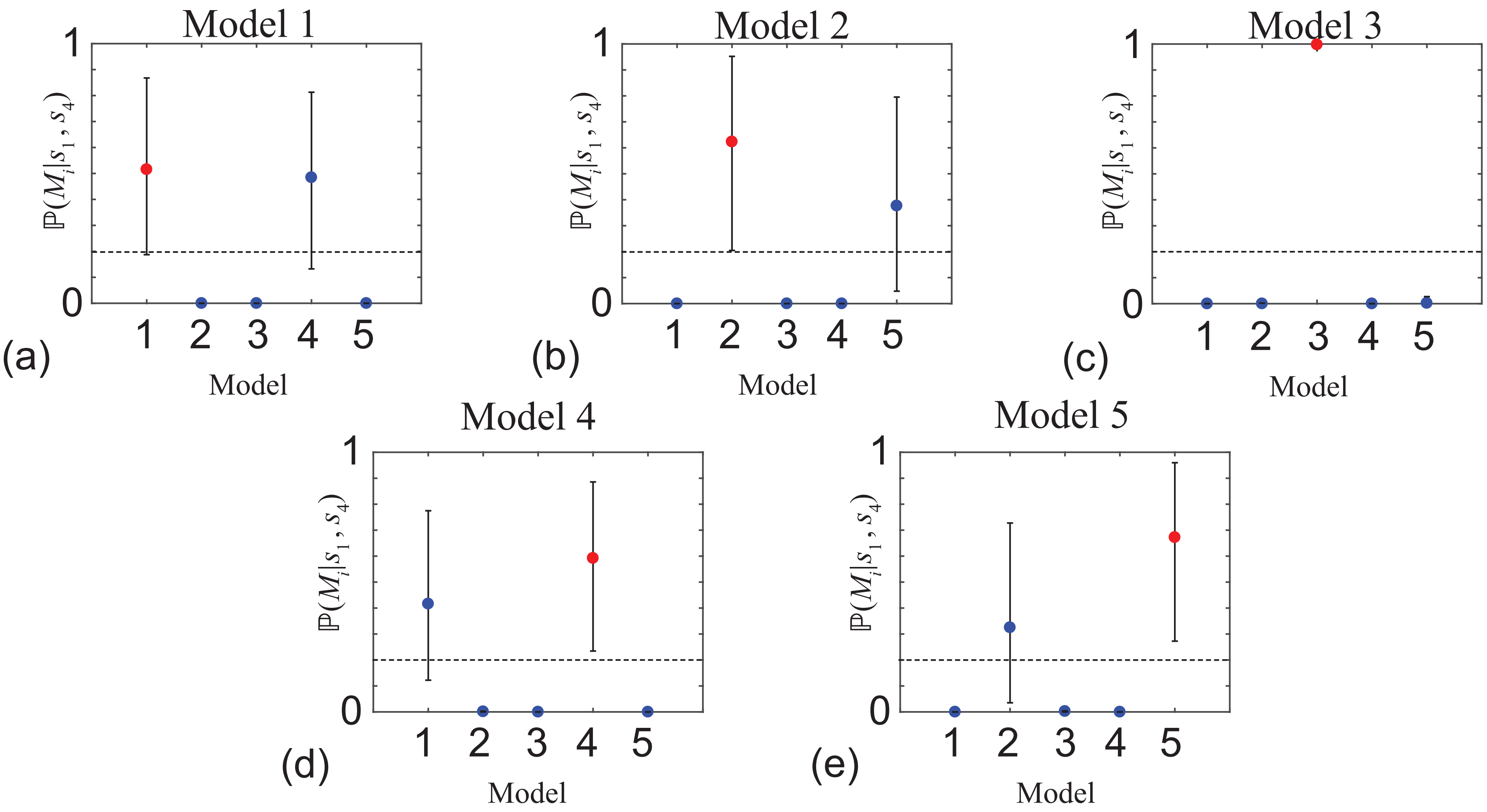


Figure 5: **Model identification with the bivariate summary statistic, $(s_1, s_2)$.** Results in (a)-(e) show point estimates of $\mathbb{P}(M_i|s)$ for $i = 1, 2, \ldots, 5$ and the uncertainty in the estimate is indicated by the error bars. The point estimates correspond to the sample mean and the error bar corresponds to the sample mean plus or minus one sample standard deviation, both calculated using 1000 identically prepared estimates of $\mathbb{P}(M_i|s)$ for $i = 1, 2, \ldots, 5$ with $J = 5$. Each subfigure shows a horizontal line at 1/5, indicating the prior distribution of $\mathbb{P}(M_i)$ for $i = 1, 2, \ldots, 5$. In each subfigure, maximum value of $\max_{i=1,2,\ldots 5}[\mathbb{P}(M_i|s)]$ is shown in red, and in each case this is the correct model choice. All results correspond to $\lambda = 0.05$ /h.

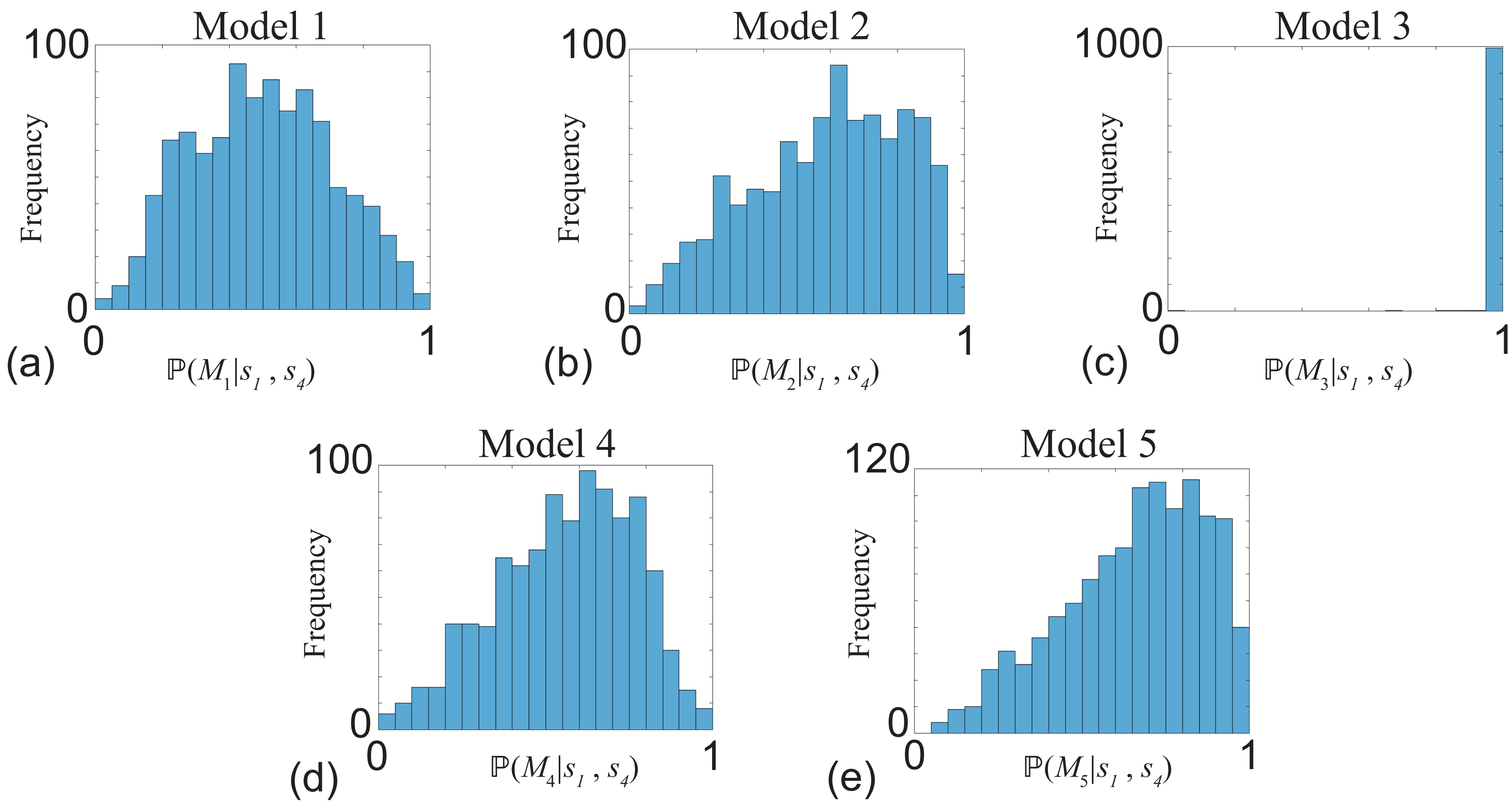


Figure 6: **Distribution of $\mathbb{P}(M_i|s)$ using the bivariate summary statistic, $(s_1, s_4)$.** Results in (a)-(e) show distributions of estimates of $\mathbb{P}(M_i|s)$ calculated using 1000 identically prepared estimates of $\mathbb{P}(M_i|s)$ for $i = 1, 2, \ldots, 5$ with $J = 5$. All results correspond to $\lambda = 0.05$ /h.
